# Synergism and antagonism of two distinct, but confused, Nrf1 factors in integral regulation of the nuclear-to-mitochondrial respiratory and antioxidant transcription networks

**DOI:** 10.1101/2020.02.12.945568

**Authors:** Shuwei Zhang, Yangxu Deng, Yuancai Xiang, Shaofan Hu, Lu Qiu, Yiguo Zhang

**Author notes:** Contributed equally to this work (Shuwei Zhang, Yangxu Deng). Correspondence should be addressed to Yiguo Zhang, or).

## Abstract

There is hitherto no literature available for explaining two distinct, but confused Nrf1 transcription factors, because they shared the same abbreviations from nuclear factor erythroid 2-related factor 1 (also called Nfe2l1) and nuclear respiratory factor (originally designated α-Pal). Thus, we have here identified that Nfe2l1^Nrf1^ and α-Pal^NRF1^ exert synergistic and antagonistic roles in integrative regulation of the nuclear-to-mitochondrial respiratory and antioxidant transcription profiles. In mouse embryonic fibroblasts (MEFs), knockout of *Nfe2l1*^*–/–*^ leads to substantial decreases in expression levels of α-Pal^NRF1^ and Nfe2l2, together with TFAM (mitochondrial transcription factor A) and other target genes. Similar inhibitory results were determined in *Nfe2l2*^*–/–*^ MEFs, with an exception that *GSTa1* and *Aldh1a1* were distinguishably up-regulated in *Nfe2l1*^*–/–*^ MEFs. Such synergistic contributions of Nfe2l1 and Nfe2l2 to the positive regulation of α-Pal^NRF1^ and TFAM were validated in *Keap1*^*–/–*^ MEFs. However, human α-Pal^NRF1^ expression was unaltered by *hNfe2l1α*^*–/–*^, *hNfe2l2*^*–/–ΔTA*^ or even *hNfe2l1α*^*–/–*^*+siNrf2*, albeit TFAM was activated by Nfe2l1 but inhibited by Nfe2l2; such an antagonism occured in HepG2 cells. Conversely, almost all of mouse Nfe2l1, Nfe2l2 and co-target genes were down-expressed in *α-Pal*^*NRF1+/–*^ MEFs. On the contrary, up-regulation of human Nfe2l1, Nfe2l2 and relevant reporter genes took place after silencing of α-Pal^NRF1^, but their down-regulation occurred upon ectopic expression of α-Pal^NRF1^. Furtherly, Pitx2 (pituitary homeobox 2) was also identified as a direct upstream regulator of Nfe2l1 and TFAM, besides α-Pal^NRF1^. Overall, these across-talks amongst Nfe2l1, Nfe2l2 and α-Pal^NRF1^, along with Pitx2, are integrated from the endoplasmic reticulum to the nuclear-to-mitochondrial communication for targeting TFAM, in order to finely tune the cellular respiratory and antioxidant gene transcription networks, albeit they differ between the mouse and the human.

## 1. Introduction

In all life forms, distinct types of cells are the most basic unit of their biological structures and functions, because they are bounded by the membrane lipid bilayers and hence protected from extracellular oxidizing environments. Of note, eukaryotic cells are evolutionarily compartmentalized by endomembrane systems, as a unique trait that distinguishes from prokaryotic cells (1), in order to give rise to distinct functionally-specialized organelles, such as the endoplasmic reticulum (ER), mitochondria and nucleus in eukaryotic cells (2). Such great benefits can allow for cellular respiration, redox metabolism and biochemical reactions to take place in tempo-spatial order during diverse physio-pathological life processes. This is owing to the facts that such versatile organelles participate in a large number of cellular functions. Amongst them, the mitochondrion is known as a central site of cellular respiration, redox metabolism and biosynthesis, in order to meet a host of energy and growth demands (3,4).

Of particular concern is the mitochondrial origin from aerobic α-proteobacterium, which was firstly surmised to be engulfed by a primordial anaerobic eukaryote (i.e., the endosymbiont hypothesis (5)) or originally merged with its archaeal host to give rise to the first eukaryotic cell (i.e., the standpoint of comparative genomics (6,7)). Thereby, this organelle has its own genetic system with the bacteria-like features, including a compact circular mtDNA genome (from maternal inheritance), along with a simple transcription system that yields multigenic RNA transcripts, and a translational apparatus with similar antibiotic sensitivities to prokaryotic cells. During subsequent endosymbiotic evolution, most of mitochondrial original genes have been lost or transferred to the nucleus in the eukaryotic cells, so that only 37 genes are retained in mammalian mitochondria (3,5). In this genome, only 13 mitochondrial genes encode proteins as essential subunits of the respiratory chain, whilst the remaining 24 genes encode specific tRNA and rRNA required for its protein translation within the mitochondrial matrix (8,9). Such a small complement of mitochondrial genes exists over the entire evolutionary process to strike a balance between the host and its endosymbionts (10,11). However, mitochondrial genetic system only possesses a semi-autonomous nature, such that its mtDNA replication, subsequent transcription and translation are subjected to the predominant control by several nuclear gene-encoding factors (9,11-13), such as mitochondrial transcription factor A (TFAM), B1 (TFB1M) and B2 (TFB2M) (14-17)). Transcriptional expression of these three factors are regulated by nuclear respiratory factor-1 (NRF-1) (18-20). Herein, NRF-1 is also referred to as α-Pal^NRF1^, because it was originally designated α-Pal from its binding to the palindromic consensus site (5’-TGCGCATGCGCA-3’) essential for the transcription of *eIF2α* gene (21-23), which controls the general protein translation. Besides, mitochondrial structure and function, as well as its biogenesis, are monitored dominantly by nuclear genome-encoded proteins (17,24,25). Notably, all the other (except the aforementioned 13) subunits of the mitochondrial respiratory chain and oxidative phophorylation are transcriptionally controlled by nuclear respiratory factors NRF-1 (i.e., α-Pal^NRF1^) and NRF-2 (also called GA-binding protein and abbreviated GABP^NRF2^, with no homology with α-Pal^NRF1^) binding to distinct consensus sites within these nuclear-encoding respiratory gene promoters (26,27).

Apart from mitochondria, the eukaryotic endomembrane system including the ER and nucleus has been posited to originate from the bacterial-like out membrane vesicles (OMVs) released by the endosymbiont mitochondrial ancestor within the cytosol of its archaeal host at eukaryote origin (1,6). The OMV-based model accounts for i) the functional homology of the ER and mitochondrial intermembrane space with dynamic topological exchanges of Ca^2*+*^ storage and the disulfide relay system required for redox signaling; ii) the cooperative contributions of the ER and mitochondria to eukaryotic lipid synthesis, as opposed to occurring at the plasma membrane as in all prokaryotes; iii) the distinction in lipid compositions of between the endomembrane system and the plasma membrane, which originated from the fact that bacterial-like lipids replaced the archaeal lipids from the inside in the first place; and that iv) the nuclear envelope is not homologous to the plasma membrane of putative archaeal host, just due to the formation of newly-enveloped nucleus from the ER in eukaryotic cells with open mitosis (6). Recently, functional and proteomic studies have revealed the remarkable complexity of mitochondrial protein machineries with diverse functions, such as protein translocation, oxidative respiration, metabolite transport, protein quality control and the control of membrane architecture, that also interact with each other in dynamic networks (28). These protein networks form distinct membrane contact sites, for example, with the ER, that are key for the integrity of mitochondria with eukaryotic cellular functions in bioenergetic synthesis, oxidative metabolism and redox signalling. The ER–mitochondria connections can also, in turn, determine mitochondrial fusion-fission dynamics. Furtherly, the mitochondrial dynamics and inheritance during cell division and development are also tightly regulated through evolutionary conserved mechanisms(29), such that they are properly segregated and partitioned as a functional set of the organelles to each of daughter cells during cell division, oogenesis, fertilization and subsequent development, as well as to ensure the integrity of its mtDNA during genetic selection of functional genomes. Contrarily, defects in these processes can lead to cell and tissue pathologies, including cancer and other degenerative diseases (29,30). However, whether putative ER-nuclear-mitochondrial (ENUM) communication is involved in mitochondrial regulation remains elusive.

Aside from only 13 mitochondrially*-*encoded *per se* proteins (9), all other thousands of the host nucleus-encoded mitochondrial proteins for its biogenesis, structure and functions in the cellular respiration, oxidative phosphorylation, energy metabolism and redox balance are subjected to their temporospatial precision expression (10,28). Of note, a considerable number of the nucleus-encoded mitochondrial proteins are transcriptionally regulated by α-Pal^NRF1^ and GABP^NRF2^ (26,27), and also translationally monitored by α-Pal^NRF1^-targeted eIF2α (21-23). Particularly, another portion of those nucleus-encoded proteins localized within and around the outer and inner mitochondrial membranes are allowed for biosynthesis in proximity to the ribosome-budded ER, because the ER is central to biosynthesis of secretory and membrane proteins, their proper folding and processing into maturation by quality controls (31,32), before being transferred to the mitochondria and anchored within double mitochondrial membranes. Moreover, the mitochondrial intermembrane space is evolutionarily homologous to the most oxidizing ER lumen, with a lowest GSH/GSSG ratio of 1:1∼3:1, than those in the relative reducing mitochondrial matrix and nuclear environments (with a higher GSH/GSSG ratio of ∼100:1) (33-35). The reducing nucleus is also segregated by the ER-connected envelope membranes from the relative oxidizing cytoplasm (GSH/GSSH = 30:1∼50:1)(33), in which mitochondria is a major source of reactive oxygen species (ROS) produced during cellular respiration (36). However, whether the ER-derived redox signalling mechanism contributes to the nuclear-to-mitochondrial respiratory and antioxidant transcription networks remains unknown.

Amongst the putative ER-derived redox signalling machineries, the most critical antioxidant arm is mediated by the membrane-bound nuclear factor erythroid-2 p45-related factor 1 (Nrf1, also called Nfe2l1) (37,38). The abbreviation is almost identical with nuclear respiratory factor-1 (NRF-1, also called α-Pal^NRF1^), leading to the confused interpretation of relevant works, the most typical of which was published by L’Honore, *et al* (with no precision corrections, compared with its original version online in *Developmental Cell* (39)). As a matter of fact, Nfe2l1 (i.e. Nfe2l1^Nrf1^) is significantly distinctive from α-Pal^NRF1^, because the former Nrf1 belongs to the cap’n’collar (CNC) basic region-lucine zipper (bZIP) family that regulate distinct subsets of antioxidant/electrophilite response element (ARE/EpRE, 5’-TGAC/GnnnGC-3’)-driven cytoprotective genes (37,40). Besides, this conserved CNC-bZIP family also includes Nfe2l2^Nrf2^ that is negatively regulated by Kelch-like ECH-associated protein 1 (Keap1, which is an adaptor subunit of Cullin 3-based E3 ubiquitin ligase and also is a key sensor for oxidative and electrophilic stresses) (40). However, hitherto no literature is available for explaining two distinct, but confused Nrf1 transcription factors, albeit they shared the same abbreviations. Herein, we have identified that Nfe2l1^Nrf1^ and α-Pal^NRF1^ exert both synergistic and antagonistic roles in integrative regulation of the nuclear to mitochondrial respiratory and antioxidant transcription profiles. Knockout of *Nfe2l1*^*–/–*^ in mouse embryonic fibroblasts (MEFs) causes substantial decreases in expression levels of α-Pal^NRF1^ and Nfe2l2^Nrf2^, along with TFAM and other target genes. Similar results were obtained from *Nfe2l2*^*–/–*^ MEFs, but rather *GSTa1* and *Aldh1a1* were significantly up-regulated. Such synergistic contributions of Nfe2l1 and Nfe2l2 to the positive regulation of α-Pal^NRF1^ and TFAM were also validated in *Keap1*^*–/–*^ MEFs. By contrast, human α-Pal^NRF1^ expression was almost unaltered by *hNfe2l1α*^*–/–*^ or *hNfe2l2*^*–/–ΔTA*^, albeit its target TFAM was activated by Nfe2l1, along with antagonism against Nfe2l2 in HepG2 cells. Conversely, almost all of mouse Nfe2l1, Nfe2l2 and co-target genes were down-expressed in *α-Pal*^*NRF1+/–*^ MEFs. Further investigation revealed that human Nfe2l1, Nfe2l2 and relevant reporter genes were down-regulated by ectopic expression of α-Pal^NRF1^, but such inhibitory effects were reversed by silencing of α-Pal^NRF1^. Moreover, Pitx2 (pituitary homeobox 2) was also identified as a direct upstream regulator of Nfe2l1 and TFAM, besides α-Pal^NRF1^. Collectively, these across-talks between Nfe2l1, Nfe2l2 and α-Pal^NRF1^, together with Pitx2, are integrated from the ER-derived signalling to the nuclear-to-mitochondrial communications for their mitochondrial target TFAM, in order to finely tune distinct cellular respiratory and antioxidant gene transcription networks responsible for maintaining their subcellularly-compartmentalized organelle homeostasis, albeit they differ between the mouse and human.

## 2. Results

### 2.1 Knockout of Nfe2l1^Nrf1^ in MEFs leads to substantial decreases of Nfe2l2, α-Pal^NRF1^, TFAM and other target genes

Here, whether knockout of Nfe2l1 has an effect on the constitutive expression of Nfe2l2 and target genes in MEFs was firstly examined by real-time quantitative PCR (RT-qPCR) and Western blotting. As anticipated, loss of Nfe2l1 in mice resulted in substantial decreases in both mRNA and protein levels of Nfe2l2 expressed constitutively in *Nfe2l1*^*–/–*^ MEFs (Figure 1, A-C). Meanwhile, an according decreases in basal expression of five of the examined ARE-driven genes *HO-1* (heme oxygenase 1, HMOX1), *GCLM* (glutamate-cysteine ligase modifier subunit), *MT-1* (metallothionein 1), *GSTp* (glutathione S-transferase pi 1) and *SOD1* (superoxide dismutase 1) were determined in *Nfe2l1*^*–/–*^ MEFs (Figure 1A). By contrast, significant increases in the basal expression of *GSTa1* (glutathione S-transferase, alpha 1 (Ya)) and *Aldh1a1* (aldehyde dehydrogenase family 1, subfamily A1) were also obtained from *Nfe2l1*^*–/–*^ in MEFs (Figure 1 A), even though they were considered as Nfe2l2-target genes (41-43). Western blotting of *Nfe2l1*^*–/–*^ MEFs revealed obvious decreases in the protein abundances of HO-1, GCLM, and SOD1, as accompanied by a striking increase in Aldh1a1 protein levels (Figure 1C). In addition, it should also be noted that Nfe2l1α and its derivates were completely deleted in *Nfe2l1*^*–/–*^ MEFs, but the residual shorter isoforms ΔN, β, γ and δ were detected by Western blotting with the antibodies against amino acids 291-741 of Nfe2l1^Nrf1^ (Figure 1B)(44). Collectively, these results indicate dual bi-directional contribution of Nfe2l1 to the positive and negative regulation of ARE-driven genes by Nfe2l2 or Nfe2l1α, respectively.

**Figure 1.**
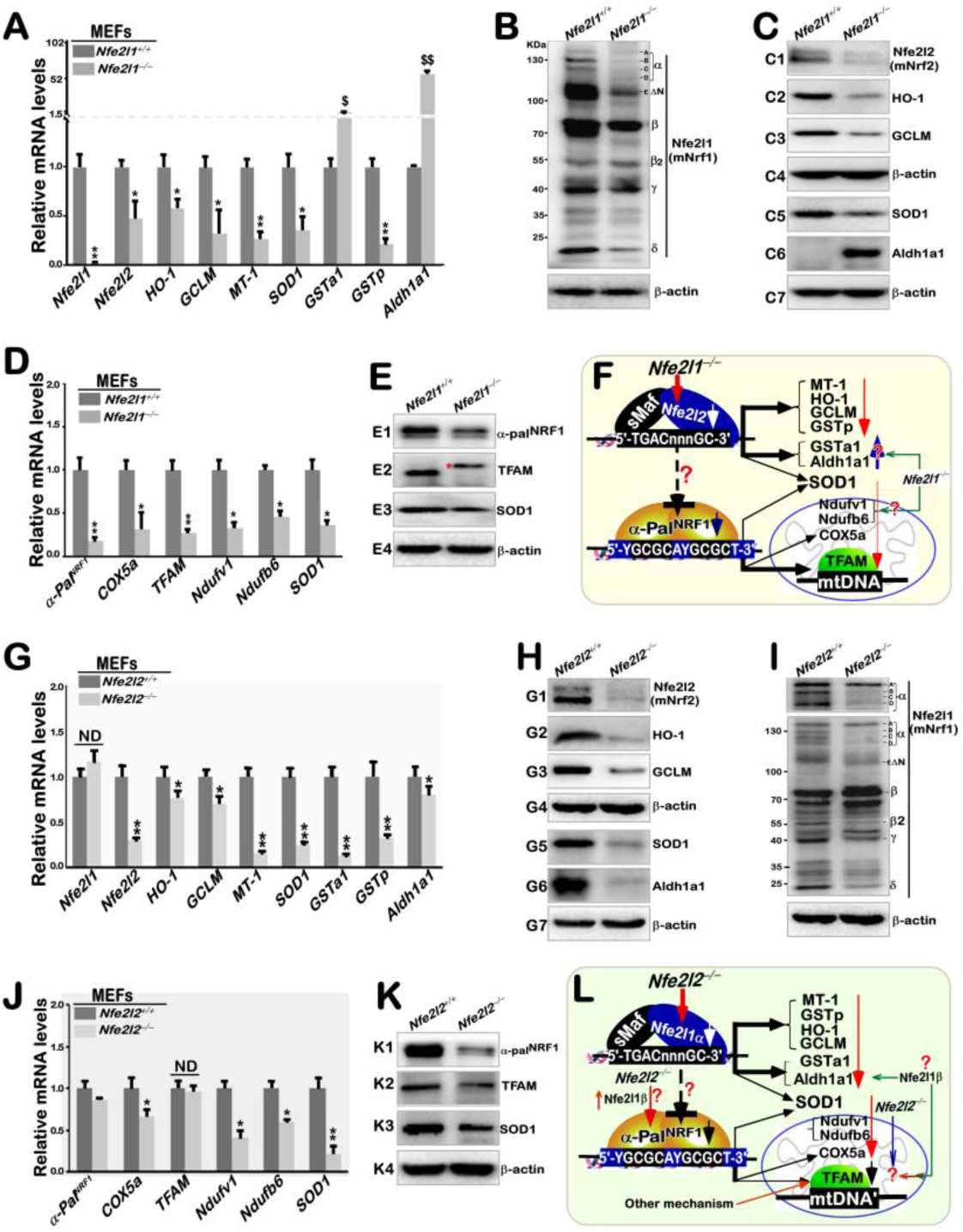
Distinct effects of *Nfe2l1^−/−^* and *Nfe2l2^−/−^* on basal expression of mouse *α-Pal^NRF1^, TFAM*, and relevant genes in MEFs. (**A**) The mRNA levels of *Nfe2l1* and *Nfe2l2*, as well as indicated antioxidant genes, were determined by real-time qPCR of *Nfe2l1*^*–/–*^ and *Nfe2l1*^*+/+*^ MEFs. The data are shown as mean ± SEM (n=3×3) with significant decreases (\**p*<0.01, \*\**p*<0.001) or increases ($, *p*<0.01, $$, *p*<0.001). (**B**) Distinct Nfe2l1 isoforms, along with a loading control β-actin, were also examined by Western blotting of *Nfe2l1*^*–/–*^ and *Nfe2l1*^*+/+*^ MEFs. (**C**) Changes in basal protein levels of Nfe2l2 and antioxidant enzymes HO-1, GCLM, SOD1 and Aldh1a1 between *Nfe2l1*^*–/–*^ and *Nfe2l1*^*+/+*^ MEFs were also observed. (**D**) Alterations in the basal mRNA expression levels of *α-pal*^*NRF1*^, *COX5a, TFAM, Ndufv1, Ndufb6 and SOD1* were unraveled by real-time qPCR of *Nfe2l1*^*–/–*^ and *Nfe2l1*^*+/+*^ MEFs. The results are shown as mean ± SEM (n=3×3) with significant decreases (\**p*<0.01, \*\**p*<0.001). (**E**) Altered proteins of α-Pal^NRF1^, TFAM and SOD1 between *Nfe2l1*^*–/–*^ and *Nfe2l1*^*+/+*^ MEFs were visualized by Western blotting. (**F**) A model is proposed to present effects of *Nfe2l1*^*–/–*^ on basal expression of *Nfe2l2, α-Pal*^*NRF1*^, *TFAM*, and relevant genes in MEFs. (**G**) Basal mRNA levels of *Nfe2l1* and Nfe2l2, as well as indicated antioxidant genes, in between *Nfe2l2*^*–/–*^ and *Nfe2l2*^*+/+*^ MEFs were also comparatively analyzed. The results are shown as mean ± SEM (n=3×3) with significant decreases (\**p*<0.01, \*\**p*<0.001) or ND (no statistical difference). (**H**) Distinct abundances of Nfe2l2 and antioxidant enzymes in between *Nfe2l2*^*–/–*^ and *Nfe2l2*^*+/+*^ MEFs were determined by Western blotting. (**I**) Altered abundances of distinct Nfe2l1 isoforms in *Nfe2l2*^*–/–*^ from *Nfe2l2*^*+/+*^ MEFs were found. (**J**) The mRNA expression levels of *α-Pal*^*NRF1*^, *COX5a, TFAM, Ndufv1, Ndufb6* and *SOD1* were analyzed by comparing real-time qPCR data from *Nfe2l2*^*–/–*^ and *Nfe2l2*^*+/+*^ MEFs. The results are shown as mean ± SEM (n=3×3) with significant decreases (\**p*<0.01, \*\**p*<0.001) or ND (no statistical difference). (**K**) Abundances of α-Pal^NRF1^, TFAM and SOD1 were compared by immunoblotting of *Nfe2l2*^*–/–*^ with *Nfe2l2*^*+/+*^ MEFs. (**L**) Another model is proposed to present potential influence of *Nfe2l2*^*–/–*^ on *Nfe2l1, α-Pal*^*NRF1*^, *TFAM*, and relevant genes (and their proteins) in MEFs.

Further RT-qPCR analysis of *Nfe2l1*^*–/–*^ MEFs revealed that loss of Nfe2l1α also led to massive decreases in mRNA expression levels of *α-Pal*^*NRF1*^ and its target genes *TFAM, COX5a* (cytochrome c oxidase subunit 5A), *Ndufv1* (NADH: ubiquinone oxidoreductase core subunit V1), *Ndufb6* (NADH:ubiquinone oxidoreductase subunit B6), and *SOD1* (Figure 1D). Substantial decreases in protein abundances of α-Pal^NRF1^ and TFAM, besides SOD1 were also visualized by Western blotting of *Nfe2l1*^*–/–*^ MEFs (Figure 1E). These demonstrate that Nfe2l1α (and its derivates) is required for expression of α-Pal^NRF1^ and its downstream targets, including TFAM (Figure 1F).

### 2.2 Knockout of Nfe2l2^Nrf2^in MEFs results in substantial decreases of α-Pal^NRF1^, TFAM and other target genes

As shown in Figure 1G, basal mRNA expression of *Nfe2l1* in MEFs was almost unaffected by knockout of *Nfe2l2*^*–/–*^, albeit all 7 of the examined Nfe2l2-targeted genes *HO-1, GCLM, MT-1, SOD1, GSTa1, GSTp*, and *Aldh1a1* were down-regulated by loss of *Nfe2l2* to different lower extents, when compared to their equivalent levels measured in wild-type (*Nfe2l2*^*+/+*^) cells. The latter notion is corroborate by evidence showing significant decreases in the constitutive protein expression of HO-1, GCLM, SOD1 and Aldh1a1 (Figure 1H). interestingly, Western blotting revealed that the full-length Nfe2l1α abundance was only marginally reduced in *Nfe2l2*^*–/–*^ MEFs, but its processed isoforms of protein-B to -D were largely abolished by *Nfe2l2*^*–/–*^, also as accompanied by a remarkable increased abundance of Nfe2l1β (Figure 1I). These suggest that possibly proteolytic processing of Nfe2l1α to yield distinct isoforms may be influenced by Nfe2l2.

Comparative analysis of *Nfe2l2*^*–/–*^ and wild-type MEFs unraveled almost no changes in mRNA expression levels of α*-Pal*^*NRF1*^ and *TFAM* (Figure 1J). However, both protein levels of α-Pal^NRF1^ and TFAM were markedly down-regulated by loss of Nfe2l2 (Figure 1K), implying that their stabilization may be monitored by Nfe2l2 (Figure 1L). Intriguingly, mRNA expression of *COX5a, Ndufv1* and *Ndufb6*, besides *SOD1*, were down-regulated to varying degrees in *Nfe2l2*^*–/–*^ MEFs (Figure 1J). This suggests that Nfe2l2 is likely required for controlling the basal expression of these three mitochondrial respiratory genes regulated by an α-Pal^NRF1^-independent mechanism (Figure 1L).

### 2.3 Significant increases of Nfe2l1, Nfe2l2, α-Pal^NRF1^, TFAM and other target genes in Keap1^−/−^MEFs

Knockout of the Nfe2l2-inhibitor Keap1 caused significant increases in protein abundances of Nfe2l2 and its down-stream targets HO-1, GCLM, Aldh1a1 and SOD1 in *Keap1*^*–/–*^ MEFs (Figure 2A), as consistent with the previous work (45). Of note, basal abundances of Nfe2l1-α, ΔN, γ and δ, but not -β, isoforms were incremented in *Keap1*^*–/–*^ MEFs (Figure 2B), albeit its mRNA expression was unaltered by loss of Keap1 (Figure 2C). However, a substantial decrease of *Nfe2l2* at its basal mRNA expression levels was determined in *Keap1*^*–/–*^ MEFs (Figure 2C). These observations demonstrate that increased abundances of Nfe2l2, and also possibly Nfe2l1, proteins are negatively monitored by Keap1-mediated degradation. Such co-activation of Nfe2l2 and Nfe2l1 by *Keap1*^*–/–*^ led to distinct extents of up-regulation of ARE-driven genes *HO-1, GSTa1, Aldh1a1, GCLM* and *SOD1*, but not *GSTp* or *MT-1* (Figure 2C). In fact, *GSTp* was unaffected, whilst *MT-1* was marginally suppressed, by *Keap1*^*–/–*^ *-*led co-active Nfe2l2 and Nfe2l1 (Figure 2C), even though *GSTp* and *MT-1* were substantially down-regulated in either *Nfe2l1*^*–/–*^ or *Nfe2l2*^*–/–*^ MEFs (Figure 1, A & G), no matter whether Nfe2l1β was decreased in *Nfe2l1*^*–/–*^ (Figure 1B), but conversely increased in *Nfe2l2*^*–/–*^ MEFs (Figure 1H). Thereby, it is inferable that both *GSTp* and *MT-1* genes (and possibly *Nfe2l2 per se*) are also likely subjected to the putative dominant negative regulation by, at least, a not-yet-identified transcription factor competitively against the positive effects of Nfe2l2 and Nfe2l1 in *Keap1*^*–/–*^ MEFs, as described by Hayes, *et al* (46)_º_

**Figure 2.**
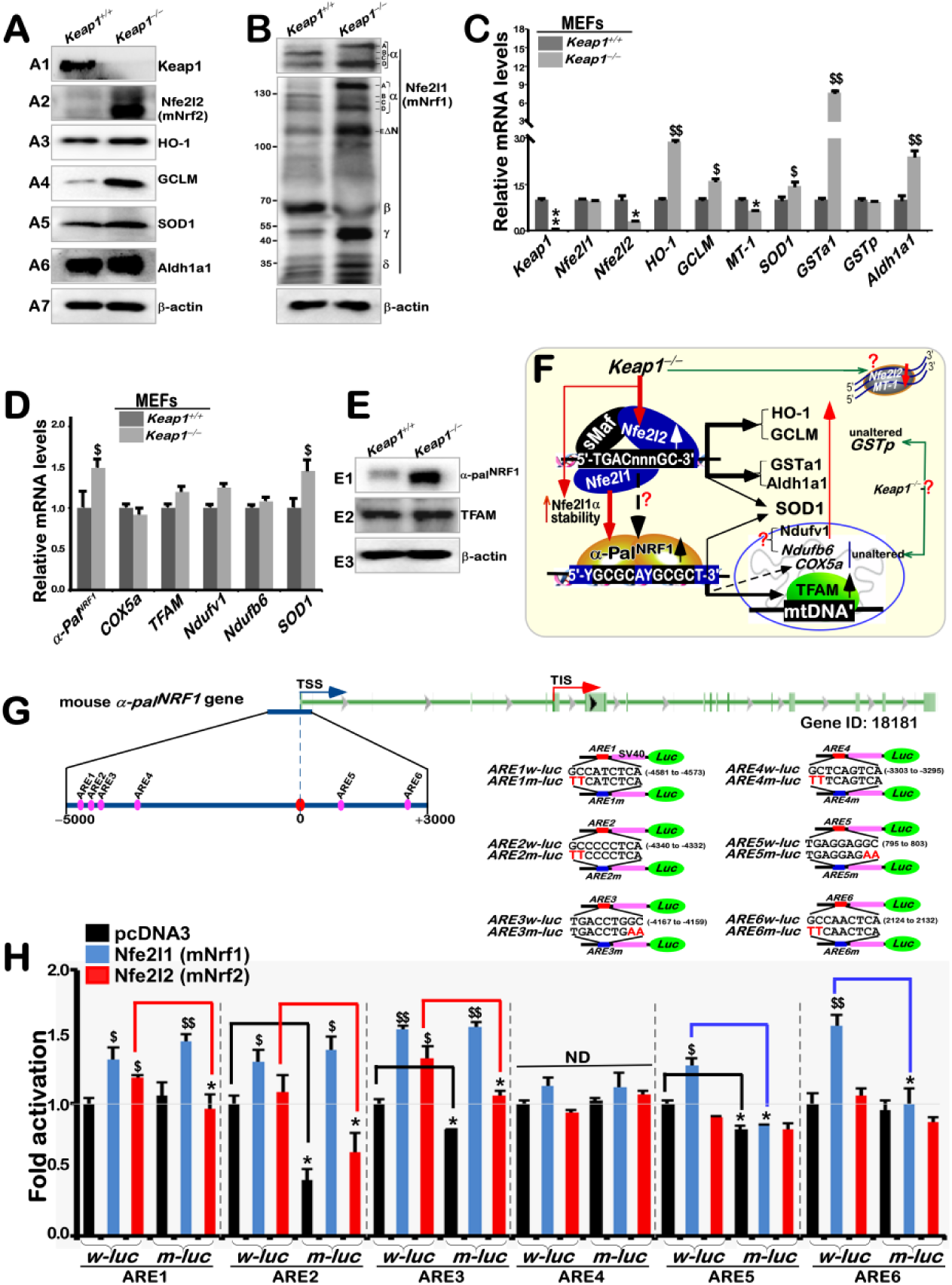
Distinct roles of key redox control genes in basal expression of mouse *α-Pal^NRF1^, TFAM*, and relevant genes. **(A)** Distinct protein levels of keap1 and Nfe2l2, as well as indicated antioxidant enzymes, in between *Keap1*^*–/–*^ and *Keap1*^*+/+*^ MEFs were visualized by Western blotting. (**B**) Altered abundances of distinct Nfe2l1 isoforms between *Keap1*^*–/–*^ and *Keap1*^*+/+*^ MEFs were observed. (**C**) Basal mRNA levels of *keap1, Nfe2l1* and *Nfe2l2*, as well as indicated antioxidant genes, were determined by real-time qPCR of *Keap1*^*–/–*^ and *Keap1*^*+/+*^ MEFs. The data are shown as mean ± SEM (n = 3×3) with significant decreases (\**p*<0.01, \*\**p*<0.001) or significant increases ($, *p*<0.01, $$, *p*<0.001). (**D**) Basal mRNA levels of *α-Pal*^*NRF1*^, *COX5a, TFAM, Ndufv1, Ndufb6* and *SOD1* were also determined as described above. (**E**) Altered protein levels of α-Pal^NRF1^ and TFAM were revealed by Western blotting of *Keap1*^*–/–*^ and *Keap1*^*+/+*^ MEFs. (**F**) A model is proposed to present effects of *Keap1*^*–/–*^ on *Nfe2l1, Nfe2l2, α-Pal*^*NRF1*^, *TFAM*, and related genes (and their proteins) in MEFs. (**G)** Within the promoter region of mouse *α-Pal*^*NRF1*^ gene, the putative *ARE* sites (each with the core sequence 5’-TGAC/GnnnGC-3’) were marked (as purple dots, *left panel*). Six distinct *ARE*-driven reporters (i.e. *ARE1* to *ARE6*-*luc*) and their respective mutants were constructed into the pGL3-Promoter vector (*right panel*). (**H**) Each pair of indicated *ARE-luc* and mutants was co-transfected with the internal control pRL-TK, together with each of expression constructs for mouse Nfe2l1, Nfe2l2 or empty pcDNA3.1 into RL34 cells for 8 h, before being allowed for 24-h recovery. Subsequently, distinct *ARE*-driven luciferase activity was measured. The resultant data are shown as mean ± SEM (n=3×3) with significant increases ($, *p*<0.01, $$, *p*<0.001) or decreases (\**p*<0.01). ND, no statistical difference.

By contrast, further examination of *Keap1*^*–/–*^ MEFs uncovered remarkable increases of α-Pal^NRF1^ and TFAM in both mRNA and protein expression levels (Figure 2,D & E). This demonstrates up-regulation of both α-Pal^NRF1^ and TFAM by *Keap1*^*–/–*^ *-*led co-activation of Nfe2l2 and Nfe2l1. However, *Ndufv1* and *Ndufb6* (both encoding two distinct subunits of the mitochondrial respiratory chain complex I) were only slightly promoted, whilst expression of *COX5a* was roughly unaltered, in *Keap1*^*–/–*^ MEFs (Figure 2D). This implies a putative negative transcription factor competitively against the positive effects of α-Pal^NRF1^ on transcriptional expression of its three mitochondrially-targeted genes *Ndufv1, Ndufb6* and *COX5a* encoded in the nucleus of *Keap1*^*–/–*^ MEFs (Figure 2F).

### 2.4 Induction of ARE-driven α-Pal^NRF1^luciferase reporter genes mediated by ectopic Nfe2l1 and/or Nfe2l2

To further determine the transcriptional regulation of α-Pal^NRF1^ by Nfe2l1 and Nfe2l2, we herein created 6 pairs of the wild-type luciferase reporters driven by the indicated *ARE* consensus sites from mouse α*-Pal*^*NRF1*^promoter and their corresponding mutants (in Figure 2G). Amongst them, basal activity of ARE2, ARE3, and ARE5 was obviously prevented by their respective mutants (Figure 2H). This implies the putative regulation of these three reporters by endogenous ARE-binding factors Nfe2l1 and/or Nfe2l2. Further co-transfection of RL34 cells with an Nfe2l1^Nrf1^ expression construct revealed that transcriptional expression of the other 5 *ARE*-, except *ARE4*-, driven luciferase reporters was activated by Nfe2l1 (Figure 2H). Such Nfe2l1-mediated activity of only ARE5 and ARE6 situated within the first exon was suppressed by their mutants, but other three *ARE1, ARE2* and *ARE3* activity mediated by Nfe2l1 was largely unaffected by their mutants (Figure 2H). By contrast, transcriptional activity of *ARE1-, ARE2*- and *ARE3-luc* reporters was not only activated by Nfe2l2, but also reduced by their respective mutants to relative lower degrees. Together, these indicate that ARE1, ARE2 and ARE3 are Nfe2l2-specific regulatory sites within the mouse *α-Pal*^*NRF1*^ promoter region, whilst ARE5 and ARE6 are Nfe2l1-specific regulatory sites within the 5’-noncoding exon one-encoded region of *α-Pal*^*NRF1*^. In addition, *ARE1-, ARE2*- and *ARE3-*mutant reporters were also activated by Nfe2l1, but not by Nfe2l2. This suggests a possibility that the three ARE mutants from the *α-Pal*^*NRF1*^ promoter region could also serve as artificial consensus sites recruiting Nfe2l1-downstream transcription factors. Thus, it is inferable that up-regulation of *ARE1, ARE2* and *ARE3* activity to mediate *α-Pal*^*NRF*^ expression may also be mediated by Nfe2l1 directly and/or indirectly its target downstream factors.

### 2.5 Down-regulation of mouse Nfe2l1, Nfe2l2 and ARE-driven genes by heterogeneous deletion of α-Pal^NRF1^ in MEFs

In turn, to explore whether α-Pal^NRF1^ contributes to *Nfe2l1, Nfe2l2 and ARE-*driven gene expression, we employed the CRISPR-Cas9-mediated genome-editing of wild-type MEFs to create a cell line called *α-Pal*^*NRF1+/–*^. This cell line was further confirmed to be true by its DNA sequencing and RT-qPCR (Figures 3A & S1). Accordingly, basal expression levels of α-Pal^NRF1^-target genes *TFAM, Ndufv1, Ndufb6, COX5a* and *SOD1* were markedly decreased, as α-Pal^NRF1^ was deleted heterogeneously in *α-Pal*^*NRF1+/–*^ MEFs (Figures 3, A & B). Further RT-qPCR analysis revealed that expression of *Nfe2l1, Nfe2l2* and *ARE-*driven genes *GCLM, MT-1, Aldhqa1, GSTa1 and GSTp1*, besides *SOD1*, were reduced by *α-Pal*^*NRF1+/–*^ to different lower extents, when compared with those equivalents of *α-Pal*^*NRF1+/+*^ cells (Figure 3C). Western blotting of *α-Pal*^*NRF1+/–*^ cells unraveled substantial decreases in distinct isoforms of mouse *Nfe2l1* and *Nfe2l2* proteins (Figure 3, D & E). Consistently, abundances of HO-1, GCLM, Aldhqa1 and SOD1 proteins were strikingly decreased in *α-Pal*^*NRF1+/–*^ MEFs (Figure 3E). Collectively, these findings demonstrate a putative contribution of α-Pal^NRF1^ (and its signalling) to expression of *Nfe2l1, Nfe2l2* and *ARE-*driven target genes (Figure 3F). In addition to their transcriptional expression, their post-translational processes may also be influenced by α-Pal^NRF1^ signalling. This is based on the observations that significantly decreases in abundances of Nfe2l1 and HO-1 proteins were, rather, accompanied by less or no changes in their mRNA expression levels, by comparison of *α-Pal*^*NRF1+/–*^ with wild-type (*α-Pal*^*NRF1+/+*^) MEFs (Figure 3, C-E).

**Figure 3.**
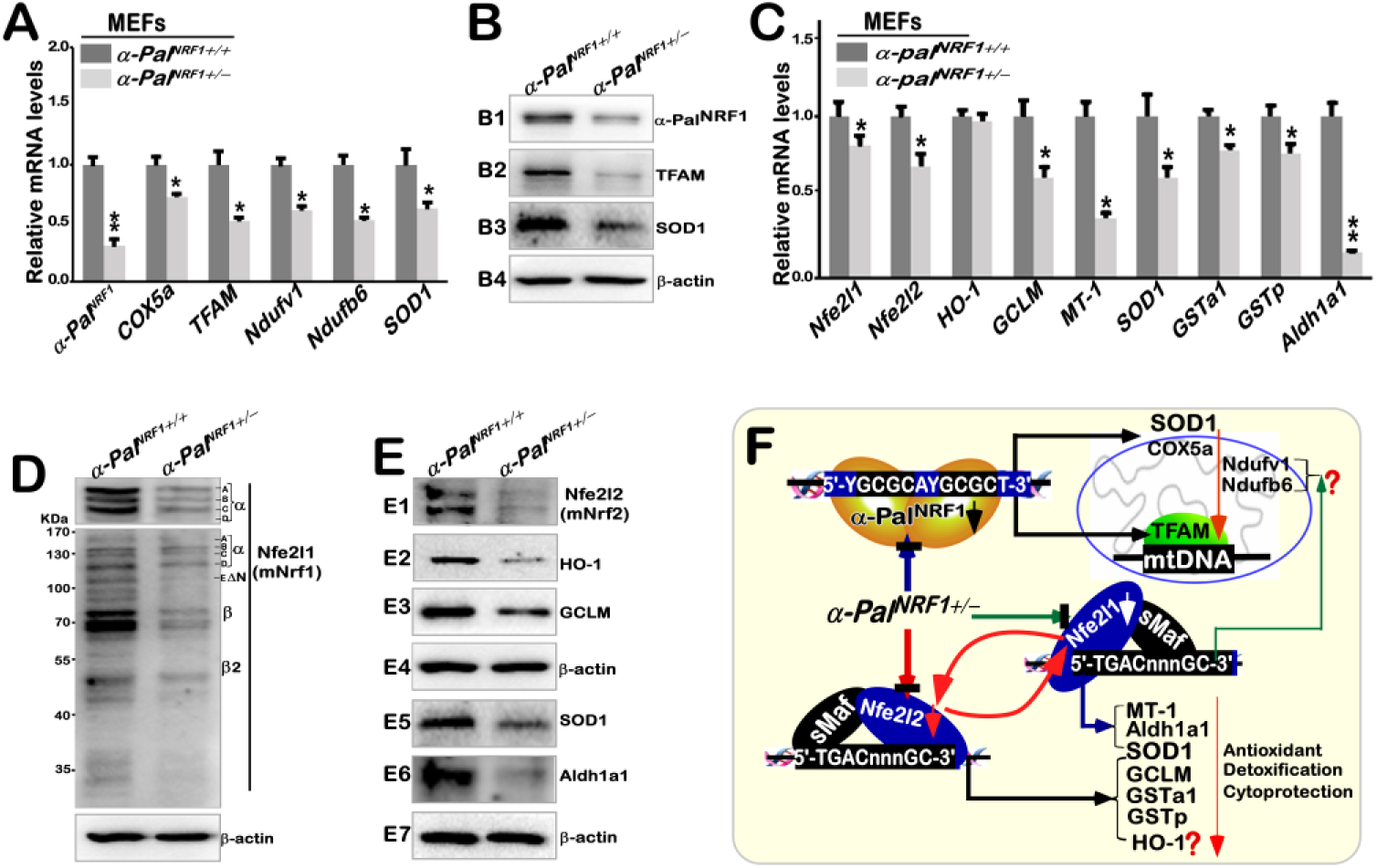
Distinct effects of mouse *α-pal^NRF1+/–^* on the expression of *Nfe2l1, Nfe2l2, TFAM and* related genes. (**A**) Altered mRNA levels of *α-pal*^*NRF1*^, *COX5a, TFAM, Ndufv1, Ndufb6* and *SOD1* in *α-pal*^*NRF1+/–*^ MEFs were compared with wild-type (*α-pal*^*NRF1+/+*^). The data are shown as mean ± SEM (n=3×3) with significant decreases (\**p*<0.01, \*\**p*<0.001). (**B**) Significant changes inα-Pal^NRF1^, TFAM and SOD1 proteins were detected by Western blotting of *α-pal*^*NRF1+/–*^ and *α-pal*^*NRF1+/+*^ MEFs. (**C**) Changes in basal mRNA levels of *Nfe2l1* and *Nfe2l2*, as well as indicated antioxidant genes, were determined by real-time qPCR of *α-pal*^*NRF1+/–*^ MEFs, as compared with *α-pal*^*NRF1+/+*^ MEFs. The results are shown as mean ± SEM (n=3×3) with significant decreases (\**p*<0.01, \*\**p*<0.001). (**D**) Altered abundances of distinct Nfe2l1 isoforms in between *α-pal*^*NRF1+/–*^ and *α-pal*^*NRF1+/+*^ MEFs were also visualized. (**E**) Altered protein levels of Nfe2l2 and indicated antioxidant enzymes (HO-1, GCLM, SOD1 and Aldh1a1) were further unraveled by Western blotting of *α-pal*^*NRF1+/–*^ and *α-pal*^*NRF1+/+*^ MEFs. (**F**) A model is proposed for distinct effects of *α-pal*^*NRF1+/–*^ on the nuclear-to-mitochondrial respiratory and antioxidant genes in MEFs.

### 2.6 No alterations of α-Pal^NRF1^ in hNfe2l1α^−/−^ or hNfe2l2^−/−ΔTA^ are accompanied by opposing changes of TFAM

Next, we examined the inter-regulatory effects of between human Nfe2l1 and Nfe2l2 on α-Pal^NRF1^, TFAM and other target genes. Unexpectedly, almost no changes in protein and mRNA expression levels of α-Pal^NRF1^ were determined in either *hNfe2l1*α^*–/–*^ or *hNfe2l2*^*–/–Δ****TA***^ cells (both lines were established by Qiu, *et al* (38,47)), when compared with those equivalents obtained from wild-type cells (Figures 4, A6 & B). Accordingly, no or less alterations in basal expression of *Ndufv1, Ndufb6* and *COX5a* by *hNfe2l1*α^*–/–*^ or *hNfe2l2*^*–/–Δ****TA***^ were also observed (Figure 4B). However, α-Pal^NRF1^-target TFAM at its protein and mRNA levels was strikingly down-regulated in *hNfe2l1*α^*–/–*^ cells, but rather significantly up-regulated in *hNfe2l2*^*–/–Δ****TA***^ cells, by comparison with their wild-type controls (Figures 4, A7 & B). These observations were further substantiated by transcriptomic sequencing (Figure 4C). Collectively, these results indicate that both Nfe2l1 and Nfe2l2 contributes respectively to the putative positive and negative regulation of human TFAM expression, possibly through a mechanism independent of α-Pal^NRF1^. This notion is supported by the evidence showing that a substantial increase in protein abundances of TFAM, rather than α-Pal^NRF1^, was resulted from over-expression of ectopic Nfe2l1 (i.e., Nrf1-V5, in Figure 4, *cf*. D3 with D4).

**Figure 4.**
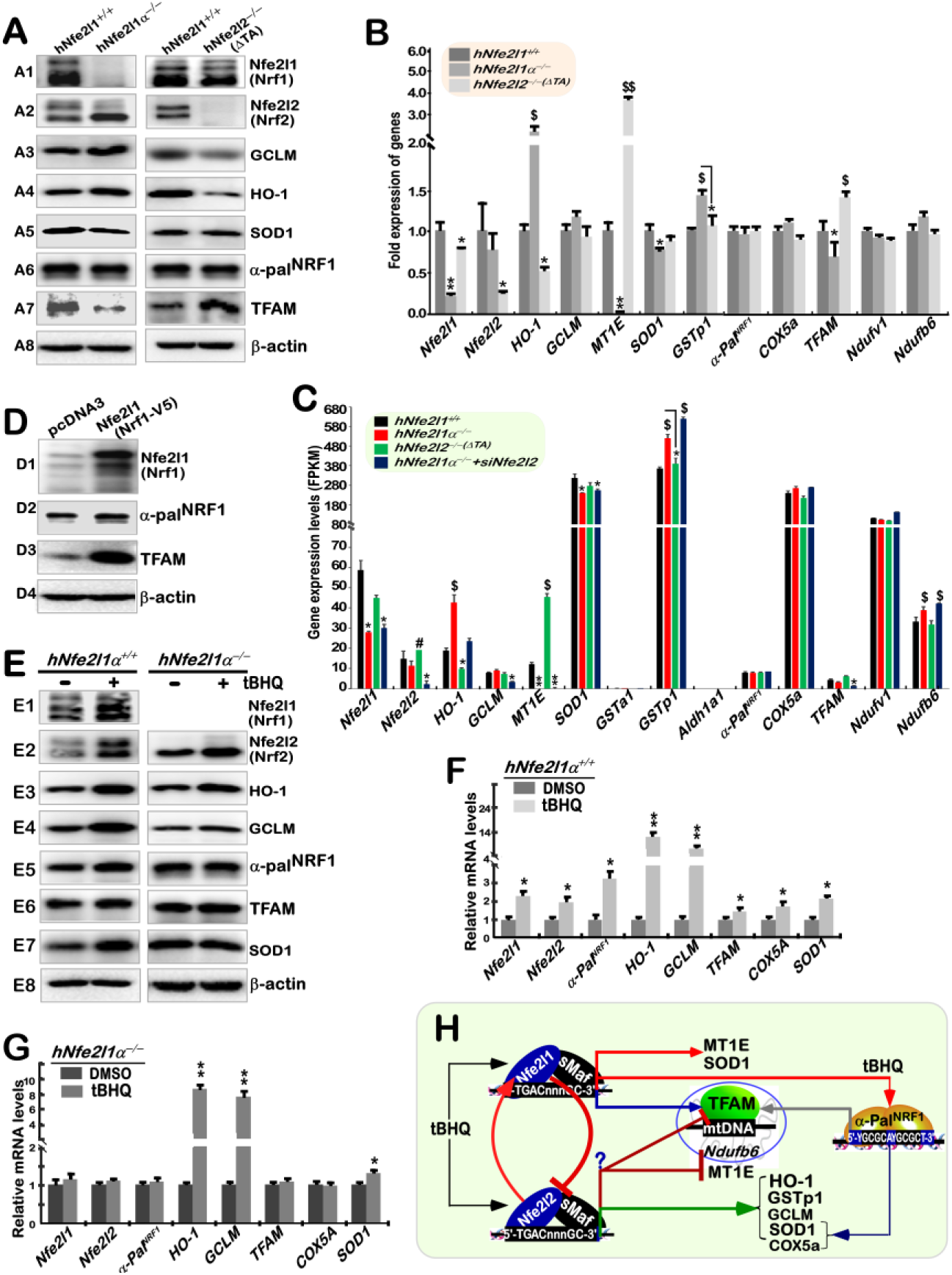
Distinct changes of human α-Pal^NRF1^ and TFAM in HepG2-derived *hNfe2l1α^−/−^* and *hNfe2l2^−/−^* cell lines. (**A**) Distinct protein levels of human Nfe2l1, Nfe2l2, α-Pal^NRF1^, TFAM, as well as other relevant proteins were determined by Western blotting of *hNfe2l1α^−/−^, hNfe2l2*^*–/–*^ and wild-type HepG2 cells. (**B**) Basal mRNA expression levels of *Nfe2l1, Nfe2l2, α-Pal*^*NRF1*^, *TFAM*, and other indicated genes were examined by real-time qPCR of *hNfe2l1α^−/−^, hNfe2l2*^*–/–*^ and wild-type HepG2 cells. The resultant data are shown as mean ± SEM (n=3×3) with significant decreases (\**p*<0.01, \*\**p*<0.001) or significant increases ($, *p*<0.01, $$, *p*<0.001). (**C**) The FPKM (Reads Per Kilobase per Million mapped reads) value of *Nfe2l1, Nfe2l2, α-Pal*^*NRF1*^, *TFAM*, and other indicated genes were obtained by RNA-Sequencing of *hNfe2l1α^−/−^, hNfe2l2*^*–/–*^, *hNfe2l1α^−/−^+siNfe2l2* and *hNfe2l1/2*^*+/+*^. (**D**) Different protein levels of Nfe2l1, α-Pal^NRF1^ and TFAM were examined in HepG2 cells that had been transfected with a hNfe2l1 expression construct or empty pcDNA3.1. (**E**) Distinct inducible alterations in abundances of Nfe2l1, Nfe2l2, α-Pal^NRF1^, TFAM, HO-1, GCLM and SOD1 were determined by Western blotting of either *hNfe2l1α^+/+^* or *hNfe2l1α^−/−^* that had been or not been treated with 50 μmol/L tBHQ. (**F, G**) Distinct inducible mRNA levels of *Nfe2l1, Nfe2l2, α-Pal*^*NRF1*^, *HO-1, GCLM, TFAM, COX5a* and *SOD1* were revealed by real-time qPCR of both tBHQ-stimulated lines of *hNfe2l1α^+/+^* cells (*F*) and *hNfe2l1α^−/−^* cells (*G*). The resultant data are shown as mean ± SEM (n=3×3) with significant increases (\**p*<0.01, \*\**p*<0.001). (**H**) A model is assumed to present cross-talks between human Nfe2l1 and Nfe2l2, along with distinct effects on human α-Pal^NRF1^, TFAM and other gene expression, particularly upon stimulation by tBHQ.

In addition, the inter-regulatory effects of between human Nfe2l1 and Nfe2l2 on the ARE-driven genes were also validated, as consistent with our previous reports (38,47). Of note, knockout of *hNfe2l1*α^*–/–*^ caused an marked increase in human Nfe2l2 protein abundances, but not its mRNA expression levels (Figure 4, A2, B and C). In turn, *hNfe2l2*^*–/–Δ****TA***^ cells gave rise to an obvious reduction of post-synthetically-processed Nfe2l1 isoforms and its mRNA levels (Figures 4, A1, B and C).

Western blotting of *hNfe2l1*α^*–/–*^ revealed that massive increases of both GCLM and HO-1 proteins were coincided with hyper-active Nfe2l2 (Figure 4, *A3 & A4*). RT-qPCR analysis unraveled that mRNA expression of *HO-1* was substantially up-regulated in *hNfe2l1*α^*–/–*^ cells, but also significantly down-regulated in *hNfe2l2*^*–/–Δ****TA***^ cells (Figure 4B), whereas *GCLM* expression was roughly unaltered in these two cell lines, when compared to wild-type controls. Further transcriptomic sequencing unveiled that basal *GCLM* expression was only repressed by *hNfe2l1*α^*–/–*^*+siNrf2* (Figure 4C), implying that it is co-regulated by both Nfe2l1 and Nfe2l2. Furthermore, *GSTp1* was up-expressed in *hNfe2l1*α^*–/–*^ cells, and also further incremented in *hNfe2l1*α^*–/–*^*+siNrf2* cells, but rather unaltered in *hNfe2l2*^*–/–Δ****TA***^ cells, when compared to wild-type controls (Figures 4, B & C). By contrast, mRNA expression of *MT1E* (encoding metallothionein 1E) was almost completely abolished in *hNfe2l1*α^*–/–*^ or *hNfe2l1*α^*–/–*^*+ siNfe2l2* cells, but also enormously augmented in *hNfe2l2*^*–/–Δ****TA***^ cells (Figures 4, B & C). Overall, these demonstrate that Nfe2l1 and Nfe2l2 contributes to the positive and negative regulation of *MT1E*, respectively, on which Nfe2l1 exerts its dominant effect, but *GSTp1* and *HO-1* expression is Nfe2l2-dependent. However, SOD1 expression is Nfe2l1α-dependent, because its protein and mRNA levels were reduced in *hNfe2l1*α^*–/–*^ or *hNfe2l1*α^*–/–*^*+siNfe2l2* cells, but largely unaffected in *hNfe2l2*^*–/–Δ****TA***^ cells (Figures 4, A5, B & C).

### 2.7 Induction of α-Pal^NRF1^ and TFAM by tert-butylhydroquinone (tBHQ) is completely abolished in hNfe2l1α^−/−^ cells

Here, we determined effects of *hNfe2l1*α^*–/–*^ on tBHQ-stimulated expression of α-Pal^NRF1^ and TFAM. As anticipated, abundances of α-Pal^NRF1^, TFAM and SOD1 proteins were evidently increased following 50 μmol/L tBHQ treatment of wild-type (*hNfe2l1*α^*+/+*^) cells, when compared with the vehicle DMSO-treated controls (Figure 4, *E5 & E6*). However, similar increases of α-Pal^NRF1^, TFAM and SOD1 were not observed in *hNfe2l1*α^*–/–*^ cells, even though hyper-active Nfe2l2 was further incremented by tBHQ (*cf. E2 with E5-E7*). Subsequently, RT-qPCR analysis revealed that mRNA expression levels of *α-Pal*^*NRF1*^, *TFAM, COX5a* and *SOD1* were induced, to greater or less extents (Figure 4F), by tBHQ stimulation of Nfe2l1, but not Nfe2l2, in *hNfe2l1*α^*+/+*^ cells (Figures 4, E & F). This is based on the fact that tBHQ-inducible increases of *α-Pal*^*NRF1*^, *TFAM, COX5a*, rather than *SOD1*, appeared to be completely abolished in *hNfe2l1*α^*–/–*^ cells, albeit Nfe2l2 was hyper-expressed by tBHQ) (Figure 4G). Altogether, these demonstrate transcriptional regulation of *α-Pal*^*NRF1*^, *TFAM* and *COX5a* dominantly by tBHQ-inducible Nfe2l1, but not Nfe2l2, whilst *SOD1* expression is co-regulated by both CNC-bZIP factors (Figure 4H). In addition, tBHQ treatment of *hNfe2l1*α^*+/+*^ cells led to significant increases in protein and mRNA levels of HO-1 and GCLM (Figures 4, E3, E4 & F). Similar induction of HO-1 and GCLM by tBHQ was also obtained in *hNfe2l1*α^*–/–*^ cells (Figures 4, E3, E4 & G). Thus, it is inferable that Nfe2l2 makes a major contribute to induction of HO-1 and GCLM expression by tBHQ (Figure 4H).

### 2.8 Another contribution of α-Pal^NRF1^ to transrepression of human Nfe2l1, Nfe2l2 and ARE-driven genes

To clarify putative contributions of α-Pal^NRF1^ to transcriptional expression of *Nfe2l1*, we herein made two luciferase reporters driven by mouse and human *Nfe2l1* gene promoter regions, called *mNfe2l1-luc* and *hNfe2l1-luc*, respectively. As unexpected, ectopic α-Pal^NRF1^-V5 over-expression only caused a slight reduction in activity of *mNfe2l1-luc* reporter (Figure 5A), albeit this reporter gene was significantly induced by tBHQ (Figure 5B, *b1*). Similarly, activity of *hNfe2l1-luc* reporter was substantially activated by tBHQ (Figure 5B, *b2*), but also significantly suppressed by ectopic expression of α-Pal^NRF1^-V5 (Figure 5C, *c1 & c3*). Such over-expression of ectopic α-Pal^NRF1^-V5 also led to transcriptional repression of *GSTa2*-*ARE×6-luc* reporter (Figure 5C, *c2 & c3*). These indicate a possible contribution of α-Pal^NRF1^ to transrepression of *Nfe2l1*-*luc* and *ARE×6-luc* reporter genes.

**Figure 5.**
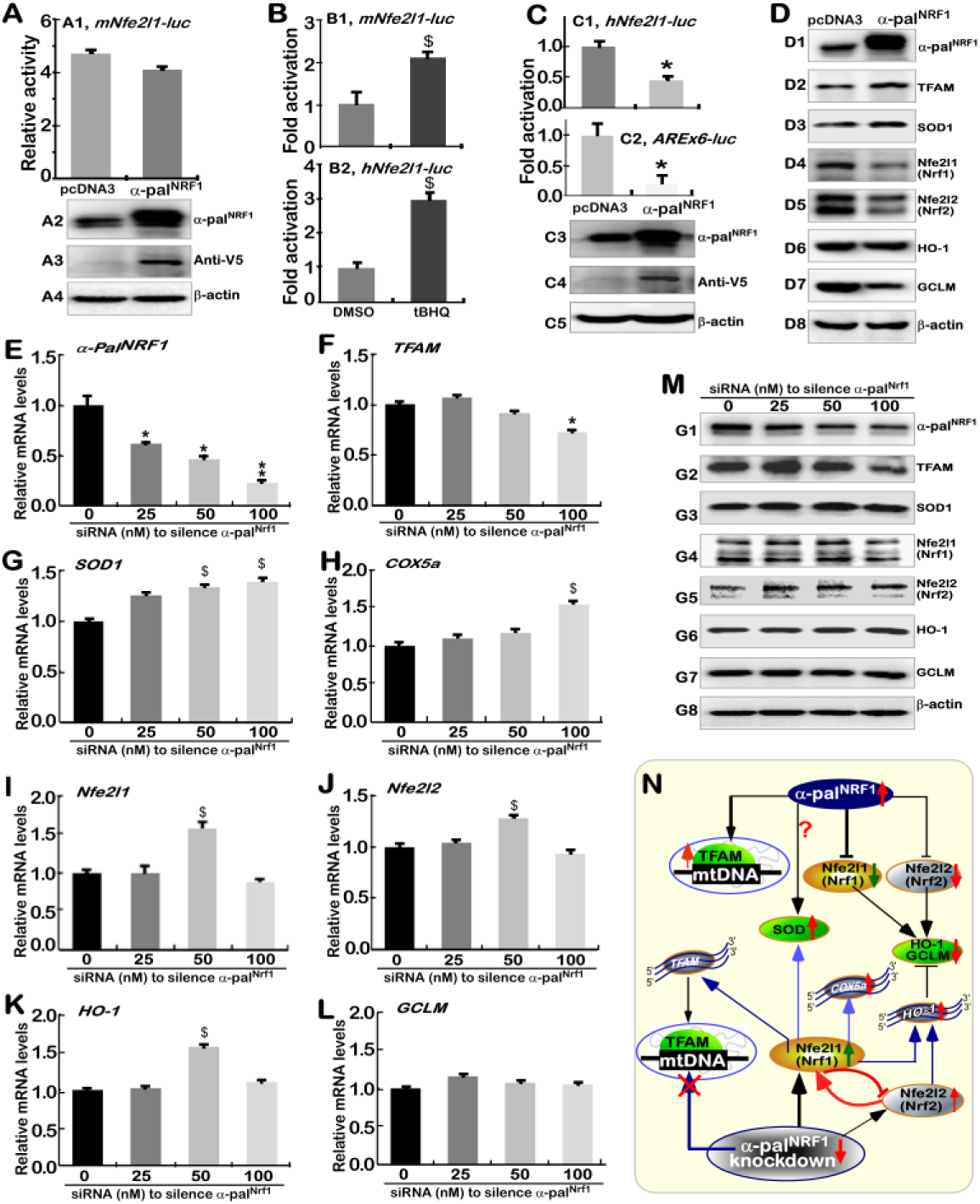
A negative effect of α-Pal^NRF1^ on *Nfe2l1, Nfe2l2* and other antioxidant genes in HepG2 cells. (**A**) HepG2 cells were co-transfected for 8 h with *mNfe2l1-luc* reporter and the pRL-TK control, along with an expression construct for mouse α-pal^NRF1^ or an empty pcDNA3.1, and then allowed for a 24-h recovery before the luciferase activity was measured (*A1*). Total lysates were also subjected to identification by Western blotting with distinct antibodies against α-Pal^NRF1^ (*A2*) or V5 tag (*A3*). (**B**) HepG2 cells were co-transfected for 8 h with *mNfe2l1-luc* (*B1*) *or hNfe2l1-luc* (*B2*), plus pRL-TK, and then treated with 50 μmol/L tBHQ or the DMSO vehicle for 24 h, before being allowed for additional 24-h recovery.Subsequently, these samples were subjected to dual luciferase assays. The results are shown as mean ± SEM (n=3×3) with significant increases ($, *p*<0.01). (**C**) HepG2 cells were co-transfected for 8 h with *hNfe2l1-luc* (*C1*) or *AREx6-luc* (*C2*) together with pRL-TK, plus a human α-Pal^NRF1^ expression construct or an empty pcDNA3.1 and then allowed for 24-h recovery from co-transfection, before the reporter activity was measured. The data are shown as mean ± SEM (n=3×3) with significant decreases (\**p*<0.01). Total cell lysates were also subjected to characterization by Western blotting with distinct antibodies against α-Pal^NRF1^ (*C3*) or V5 tag (*C4*). (**D**) Distinct protein levels of α-Pal^NRF1^, TFAM, SOD1, Nfe2l1, Nfe2l2, HO-1 and GCLM were determined by Western blotting of HepG2 cells that had been transfected with α-Pal^NRF1^ expression plasmid or an empty pcDNA3 vector. (**E**-**L**) Distinct changes in mRNA levels of *α-Pal*^*NRF1*^ (*E*), *TFAM* (*F*), *SOD1* (*G*), *COX5a* (*H*), Nfe2l1(*I*), Nfe2l2(*J*), HO-1(*K*) and GCLM(*L*) were analyzed by real-time qPCR of HepG2 cells that had been transfected with 0, 25, 50 and 100 nM of siRNA against *α-Pal*^*NRF1*^. The resulting data are shown as mean ± SEM (n=3×3) with significant decreases (\**p*<0.01, \*\**p*<0.001) or increases ($, *p*<0.01). (**M**) Such siRNA-transfected cell lysates were subjected to Western blotting analysis of distinct protein abundances as indicated. (**N**) A model is proposed to present disparate effects of α-pal^NRF1^ over-expression or its knockdown on the nuclear-to-mitochondrial respiratory and antioxidant genes in HepG2 cells.

Western blotting of α-Pal^NRF1^-expressing cells revealed that its targets TFAM and SOD1 were evidently enhanced, whilst abundances of Nfe2l1 and Nfe2l1 proteins were markedly diminished by ectopic α-Pal^NRF1^-V5 (Figure 5D, *d1-d5*). Accordingly, HO-1 and GCLM abundances were also partially inhibited by α-Pal^NRF1^-V5 (Figure 5D, *d6 & d7*). Conversely, knockdown of α-Pal^NRF1^ by its specific siRNA (Figure 5, E & M1) only caused a marginal reduction of TFAM (Figure 5, F & M2), implying coordinated transcription regulation of human TFAM by α-Pal^NRF1^ with another factor (e.g. GABP^NRF2^). By contrast, both mRNA and protein levels of SOD1 were modestly promoted by silencing of α-Pal^NRF^ (Figure 5,G & M3), implying one of other putative transcription factors competitively against knockdown of α-Pal^NRF1^. Similar results were also obtained from the case of *COX5a* (Figure 5H).

Further examinations revealed that modestly increased proteins of Nfe2l1, Nfe2l1 and HO-1 (Figure 5M, *M4-6*) were coincidently accompanied by marginal enhancements of their mRNA expression levels, particularly following transfection of HepG2 cells with 50 nM siRNA against α-Pal^NRF1^ (Figure 5, I-K). In addition, GCLM protein and mRNA levels were almost unaffected by silencing of α-Pal^NRF1^(Figure 5, L & M7). Taken together, these results indicate that α-Pal^NRF1^ contributes to the negative regulation of human Nfe2l1, Nfe2l1 and HO-1 (Figure 5N). However, no dose-responses of Nfe2l1, Nfe2l1 and HO-1 to different concentrations of α-Pal^NRF1^-silenced siRNAs indicate a possible involvement of other transcription factors (e.g., Pitx2, GABP^NRF2^). Moreover, α-Pal^NRF1^ exerts disparate effects on GCLM, possibly depending on whether Nfe2l1, Nfe2l1 and other unidentified transcription factors were also stimulated by α-Pal^NRF1^ over-expression or its RNA-silencing.

### 2.9 Identification Pitx2 as a upstream regulator to mediate transactivation of human Nfe2l1 gene

Clearly, α-Pal^NRF1^ was identified as a direct target of Pitx2/3 (pituitary homeobox 2/3, also paired like homeodomain 2/3) (39). Here, we determine whether Pitx2 also acts as a direct upstream factor to mediate transactivation of human *Nfe2l1*. As shown in Figure 6A, activity of *mNfe2l1-luc* reporter was significantly transactivated by ectopic Pitx2, when compared with the basal activity measured from its co-transfection with an empty pGL3-Basic vector. Similar results were also obtained from those *hNfe2l1-luc* reporters (Figure 6, B & C). Of note, over-expression of ectopic Pitx2 caused gradual increments in the *trans*-activity of *hNfe2l1-luc, hNfe2l1-luc1* and *hNfe2l1-luc2* reporters (Figure 6C, *C2-C4*). The *hNfe2l1-luc* activity was also strikingly induced by tBHQ to the extent of *hNfe2l1-luc1* transactivation mediated by Pitx2 (Figure 6C, *cf. C1 with C3*). In addition, four distinct consensus Pitx-response elements (PitxREs), which are situated at the upstream of the indicated promoter region to yield *hNfe2l1-luc1* (Figure 6B), were also identified here. The results revealed that only *PitxRE2-luc* activity was modestly activated by Pitx2, but basal activity of *PitxRE3-luc* and *PitxRE4-luc* was partially suppressed by Pitx2 (Figure 6D). Together, these indicate that transcriptional expression of human *Nfe2l1* gene is likely co-regulated by Pitx2 and its target factors.

**Figure 6.**
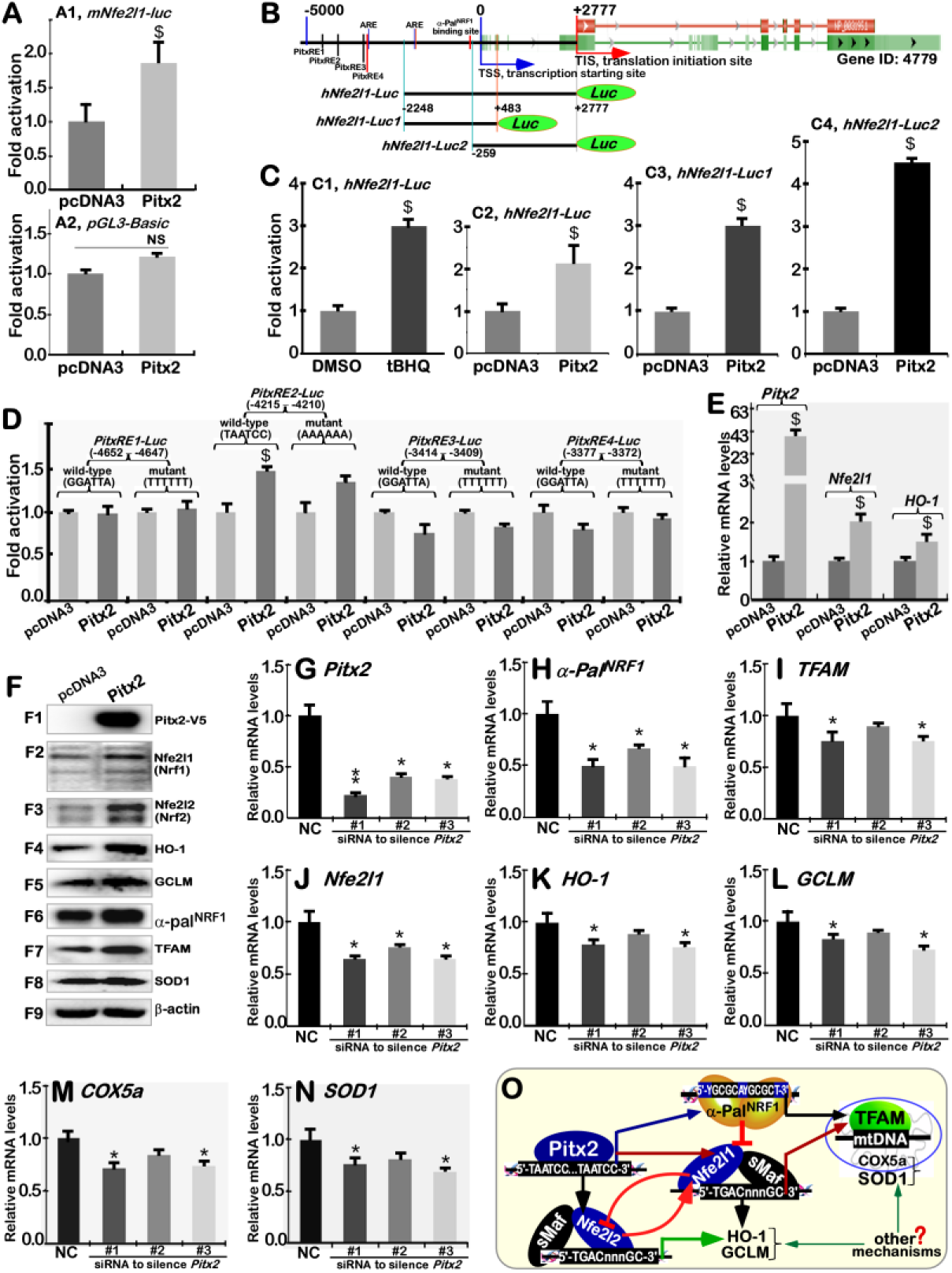
Identification of pitx2 as an upstream regulator of *Nfe2l1*, besides *α-pal^NRF1^*. (**A**) HepG2 cells were co-transfected for 8 h with *mNfe2l1-luc* (*A1*) or empty *pGL3-Basic* (*A2*), together with the pRL-TK control, plus a Pitx2 expression construct or empty pcDNA3.1, and then allowed for a 24-h recovery before the luciferase activity was measured. The data are shown as mean ± SEM (n=3×3) with significant increases ($, *p*<0.01) or NS (not significance). (**B)** Putative *cis*-regulatory binding sites for Pitx2, α-Pal^NRF1^ and AREs within the human *Nfe2l1* promoter region were indicated. Various lengths of *hNfe2l1-luc* were cloned into the pGL3-Basic vector as shown schematically. (**C**) HepG2 cells were co-transfected for 8 h (i) with *hNfe2l1-luc* and pRL-TK, and also treated for 24 h with 50 μmol/L tBHQ or the DMSO vehicle (*C1*); (ii) with *hNfe2l1-luc* (*C2*), *hNfe2l1-luc1* (*C3*), or *hNfe2l1-luc2* (*C4*), along with the pRL-TK control, plus a Pitx2 expression construct or an empty pcDNA3.1, before being allowed for additional 24-h recovery. The samples were then subjected to dual luciferase assays. The results are shown as mean ± SEM (n=3×3) with significant increases ($, *p*<0.01). (**D**) HepG2 cells were co-transfected for 8 h with each of *PitxRE-luc* reporters or their mutants, together with pRL-TK plus a Pitx2 expression construct or an empty pcDNA3.1, and then allowed for 24-h recovery. Thereafter, the reporter activity was detected and calculated as mean ± SEM (n=3×3) with significant increases ($, *p*<0.01). (**E**) Distinct mRNA levels of *Pitx2, Nfe2l1* and *HO-1* were detected by real-time qPCR of HepG2 cells that had been transfected with an expression construct for Pitx2 or an empty pcDNA3.1 vector. (**F**) Pitx2-expressing HepG2 cells were subjected to Western Blotting of Pitx2, Nfe2l1, Nfe2l2, HO-1, GCLM, α-Pal^NRF1^, TFAM and SOD1. (**G**-**N**) Distinct mRNA levels of *Pitx2* (*G*), *α-Pal*^*NRF1*^(*H*), *TFAM* (*I*), *Nfe2l1* (*I*), *HO-1* (*J*), *GCLM* (*L*), *COX5a* (*M*) and *SOD1* (*N*) were determined by real-time qPCR analysis of HepG2 cells that had been transfected with three different siRNAs against *Pitx2*. The results are shown as mean ±SEM (n=3×3) with significant decreases (\**p*<0.01, \*\**p*<0.001). (**O**) A model is proposed for potential effects of Pitx2 on Nfe2l1, Nfe2l2, α-Pal^NRF^, TFAM and other genes in HepG2 cells.

RT-qPCR analysis revealed that endogenous mRNA expression of human *Nfe2l1* and *HO-1* was increased by Pitx2 over-expression (Figure 6E). Western blotting of Pitx2-expressing cells unraveled significant increases in abundances of all 7 examined proteins Nfe2l1, Nfe2l2, HO-1, GCLM, α-Pal^NRF1^, TFAM and SOD1 (Figure 6F, *F2-F8*). These demonstrate that Pitx2 may also serve as a upstream regulator to mediate transcriptional expression of Nfe2l1 (and Nfe2l2), besides α-Pal^NRF1^. This notion is further supported by the evidence obtained from silencing of Pitx2 by its three distinct siRNA sequences (as listed in Table S1). Consequently, silencing of Pitx2 led to varying decreases in basal mRNA expression levels of *α-Pal*^*NRF1*^, *TFAM, Nfe2l1, HO-1, GCLM, COX5a* and *SOD1* to less extents than their equivalent negative controls (Figure 6, G-N). Thereby, these results unveil that Pitx2 is required for mediating transcriptional expression of *Nfe2l1, α-Pal*^*NRF1*^and their target genes (Figure 6O).

### 2.10 Identification of TFAM as another direct co-target of human Nfe2l1, Nfe2l2 and Pitx2

Although TFAM is known as a direct target of α-Pal^NRF1^ (18,48,49), we further determined whether it also serves as another direct co-target of Nfe2l1, Nfe2l2 and Pitx2. The amino acid sequences of three nuclearly-encoded TFAM from the human and mouse were aligned (as illustrated in Figure 7A), revealing that two mitochondria-targeting sequences are located within the N-terminal region of this MHG-box-binding transcription factor. Further analysis of human *TFAM* promoter showed the putative consensus *cis*-regulatory elements, such as *AREs, PitxRE* and *α-Pal*-binding site (Figure 7B). Amongst them, *ARE*- and *PitxRE*-driven luciferare reporters were established (Figure 7C). As a result, the luciferase assays of both *PitxRE-Luc* and *ARE3-Luc* activity demonstrated that transcriptional expression of TFAM is co-regulated by human Pitx2, Nfe2l1 and Nfe2l2 (Figure 7, D-F).

**Figure 7.**
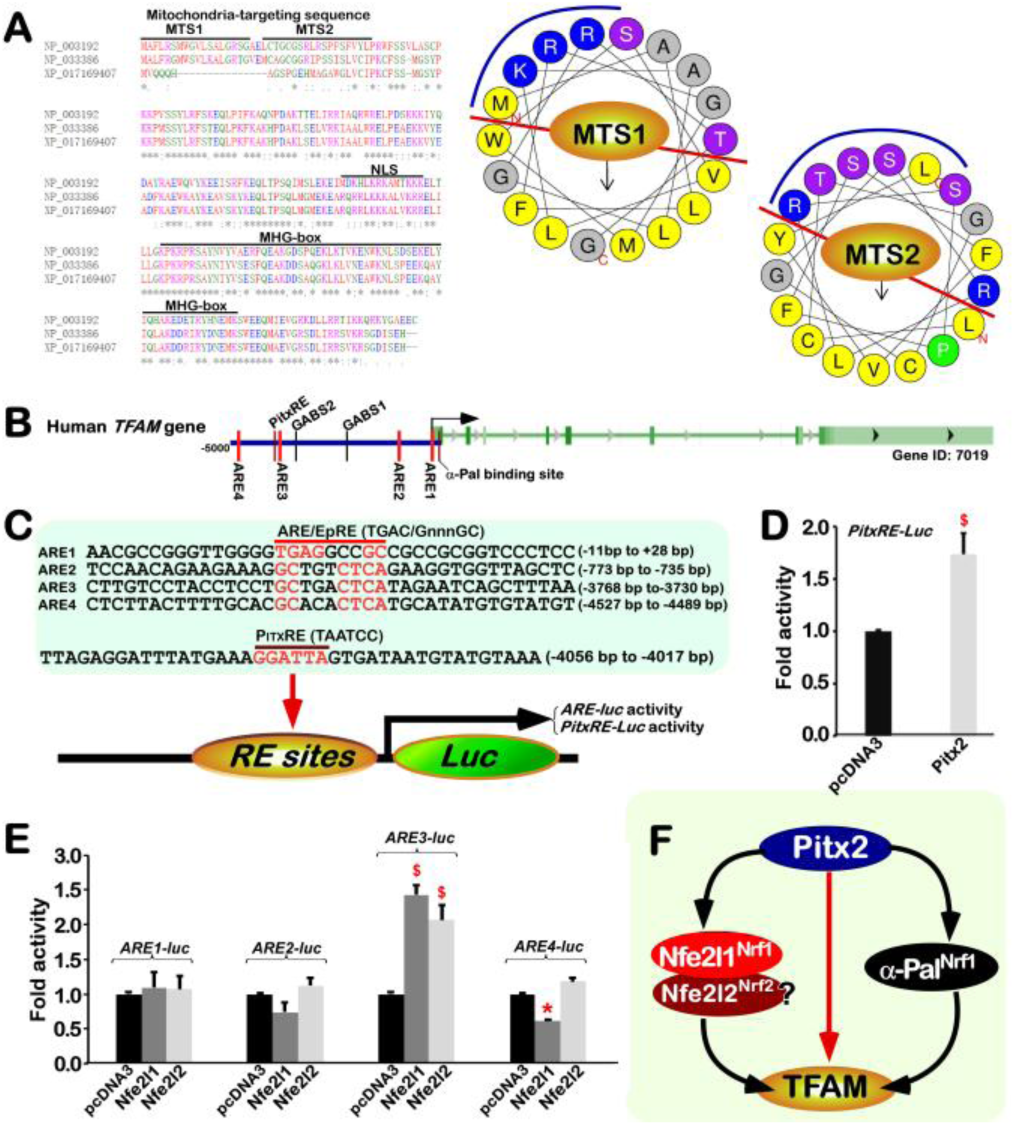
Transcriptional regulation of TFAM by Nfe2l1, Nfe2l2 and Pitx2. (**A**) Amino acid sequences of human TFAM (NP_003192) and mouse TFAMs (NP_033386 and XP_017169407) were aligned (*left panel*). Of note, mitochondria-targeting sequences (MTS), nuclear localization signal (NLS) and DNA-binding MHG-box were also indicated. Both MTS1 (aa 1-18) and MTS2 (aa 21-38) were wheeled into two similar α-helical structures. Basic arginine and lysine residues were placed on blue backgrounds, nucleophilic serine and threonine residues are on purple backgrounds, an unamiable proline residue was on a green background, and all hydrophobic amino acids were on yellow backgrounds, except for small alanine and glycine on grey backgrounds. (**B**) The putative consensus ARE sites and other cis-regulatory binding sites for Pitx2, α-Pal^NRF1^, or GABP within the human *TFAM* gene promoter region were indicated. (**C**) Four distinct *ARE*-driven (i.e. *ARE1*-*luc* to *ARE4-luc*) and another *Pitx2RE-luc*) reporters were constructed into the pGL3-Promoter vector. (**D**) HepG2 cells were co-transfected with *PitxRE-luc* and pRL-TK, plus a Pitx2 expression plasmid or an empty pcDNA3.1, and then allowed for 24-h recovery before the luciferase activity was measured. The data are shown as mean ±SEM (n=3×3) with significant increase ($, *p*<0.01). (**E**) HepG2 cells were co-transfected with each of *ARE1*-*luc* to *ARE4-luc*, together with pRL-TK plus an expression construct for Nrf1, Nrf2 or empty pcDNA3.1, and then allowed for 24-h recovery, before such *ARE*-driven activity was detected. The resultant data are shown as mean ± SEM (n=3×3) with significant increases ($, *p*<0.01) or decreases (\**p*<0.01). (**F**) A model is proposed to explain the transcriptional regulation of TFAM by Nfe2l1, Nfe2l2, Pitx2 and α-Pal^NRF1^ in HepG2 cells.

## 3. Discussion

In this study, we have, for the first time, provided a better interpretation of two totally distinctive Nrf1 transcription factors (i.e. Nfe2l1^Nrf1^ and α-Pal^NRF1^ renamed here). This is an attempt to rectify the previously-confusing explanations of both Nrf1 factors from their commonly-shared abbreviations in the literature as far as we have known(17,37,50), in which one of the most typical representative is the publication confused by L’Honore, *et al* (39). Similarly, another two Nrf2 transcription factors (51,52) are also referred to as Nfe2l2^Nrf2^ and GABPα^NRF2^, respectively. Of note, the antioxidant master Nfe2l2^Nrf2^ is evolutionarily conserved with Nfe2l1^Nrf1^, but not with the nuclear respiratory factors α-Pal^NRF1^ and GABPα^NRF2^, of which the latter two factors have also no any homology with each other. Collectively, our present study has elucidated synergistic and antagonistic roles of between Nfe2l1^Nrf1^ (and/or Nfe2l2^Nrf2^) and α-Pal^NRF1^ in integrative regulation of the signalling to the nuclear-to-mitochondrial respiratory and antioxidant transcription networks, which differ between the mouse and human (Figure 8).

**Figure 8.**
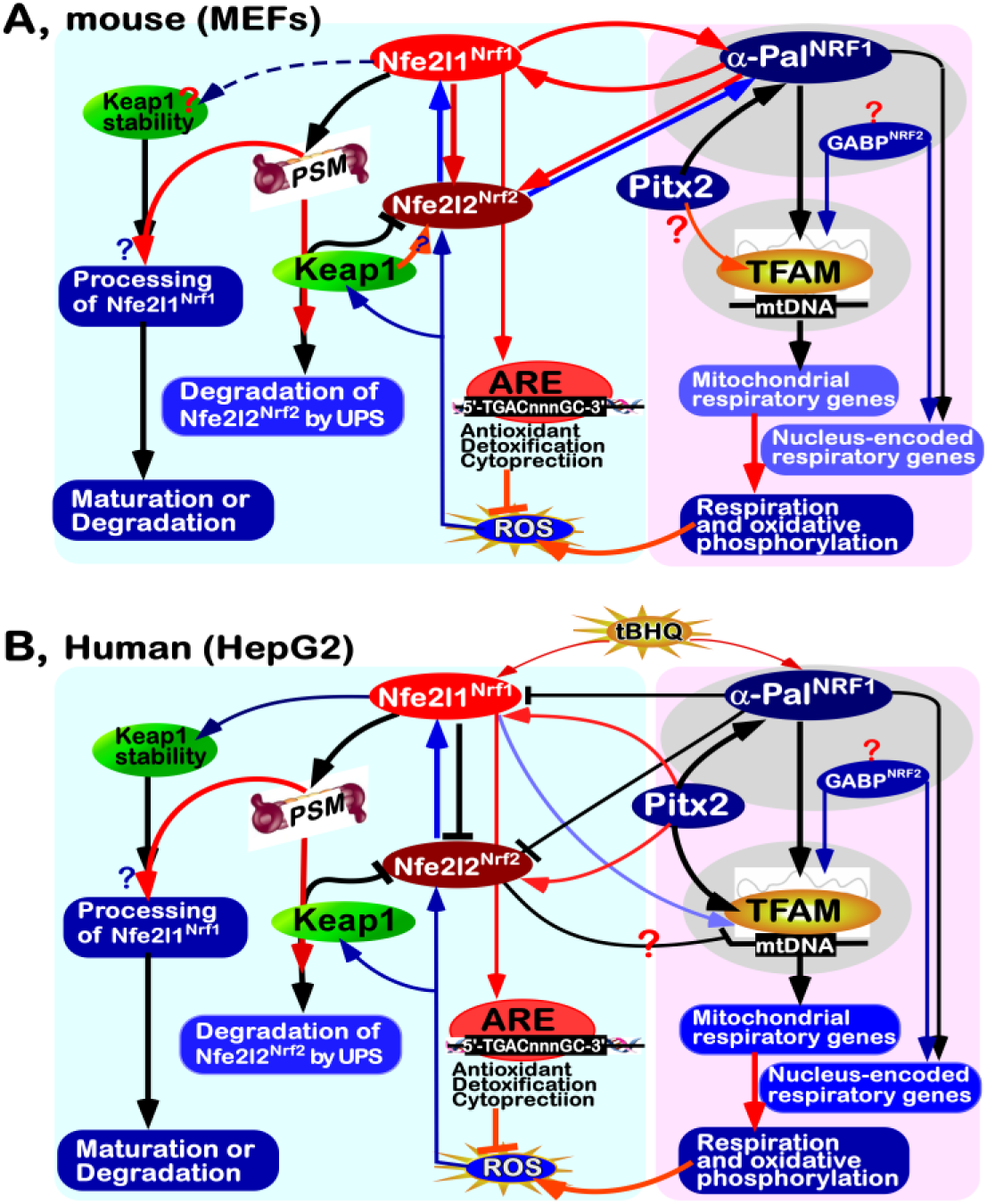
Two distinct models for integrative regulation of cellular respiratory and antioxidant transcription networks. (**A**) model is proposed on the base of experimental evidence obtained from MEFs, to give a better understanding of regulatory cross-talks between Nfe2l1^Nrf1^, Nfe2l2^Nrf2^ and α-Pal^NRF1^, along with Pitx2, which are responsible for the nuclear-to-mitochondrial respiratory and antioxidant gene transcription networks. Of note, Nfe2l1^Nrf1^ can make opposing contributions to the bi-directional regulation of Nfe2l2^Nrf2^ and α-Pal^NRF1^ by itself and target proteasome (*PSM*) at two distinct layers. Nfe2l2^Nrf2^ can also determine putative post-transcriptional regulation of Nfe2l1^Nrf1^ and α-Pal^NRF1^, but the detailed mechanisms remain unclear. In turn, α-Pal^NRF1^ also has a capability to positively regulate Nfe2l1^Nrf1^ and Nfe2l2^Nrf2^, albeit no canonic GC-rich α-Pal-binding sites within these mouse CNC-bZIP gene promoters (Table S2). Such nucleus-controlled mitochondrial respiratory and oxidative phosphorylation is also a primary source of byproduct reactive oxygen species (ROS) in cells, which can, in turn, trigger putative physiological activation of Nfe2l1^Nrf1^, Nfe2l2^Nrf2^ and α-Pal^NRF1^. Besides, GABP^NRF2^ is also required for this process, but not yet identified herein. In addition to negative regulation of Nfe2l2^Nrf2^, Keap1 can rather positively regulate transcription of this CNC-bZIP factor. (**B**) Another model is proposed, based on experimental evidence obtained from HepG2 cells, to give a better explanation of distinct cross-talks amongst Nfe2l1^Nrf1^, Nfe2l2^Nrf2^ and α-Pal^NRF1^, along with Pitx2. All these factors are converged on the nucleus-encoded mitochondria-targeting TFAM, but Nfe2l2^Nrf2^ may contribute to dual opposing effects on this mitochondrial transcription factor. Furthermore, α-Pal^NRF1^ can also make a negative contribution to transcription of human Nfe2l1^Nrf1^ and Nfe2l2^Nrf2^, albeit all three factors are activated by redox inducer tBHQ. Of note, Nfe2l2^Nrf2^ is negatively regulated by Nfe2l1^Nrf1^. In addition, human Nfe2l1^Nrf1^ is also essential for stabilization of Keap1, but whether the adaptor subunit of Cullin 3-based E3 ubiquitin ligase is involved in the proteolytic processing of Nfe2l1^Nrf1^ remains elusive.

### 3.1 Synergism of mouse Nfe2l1^Nrf1^, Nfe2l2^Nrf2^ and α-Pal^NRF1^ to coordinate the nuclear-to-mitochondrial respiratory and antioxidant gene responses in MEFs

Clearly, Nfe2l1 and Nfe2l2 are two highly-conserved members of the CNC-bZIP transcription factors predominantly regulating distinct subsets of ARE-driven cytoprotective genes against oxidative stress (37,53). These target genes are critically involved in maintaining cellular homeostasis and organ integrity during normal development and growth, as well as healthy life process, in which Nfe2l1 and Nfe2l2 cannot only exert their combinational and/or overlapping roles, and also manifest their distinctive functions. Importantly, Nfe2l1 can elicit its unique dispensable physio-pathological functions, because its loss in the mouse leads to distinct pathological phenotypes, even in the presence of Nfe2l2 (37). This implies a fact that the loss of Nfe2l1 cannot be compensated by Nfe2l2, albeit the latter Nfe2l2 has been accepted as a master regulator of antioxidant, detoxification and cytoprotective responses (53,54). Such significant distinctions in the functioning of between Nfe2l1 and Nfe2l2 are dictated by their respective intrinsic features.

Interestingly, Nfe2l1, but not Nrf2, with several domains being more highly conserved with its ancestral CNC, Skn-1 and Nach proteins (55), is located in the ER and its connected nuclear envelope membranes. Of note, there exist two extra functional domains (i.e. NTD and NST) within Nfe2l1, rather than Nfe2l2. The ER-targeting NTD of Nfe2l1 enables it to be topologically anchored within and around the membranes (56-58), whilst its NST glycodomain facilitates its proper protein folding and subsequent processing (59-61). The membrane-topology of Nfe2l1 determines its selective proteolytic processing by proteasomes and/or other cytosolic proteases in close proximity to the ER, in order to yield distinct lengths of its isoforms, before its mature CNC-bZIP factor is translocated into the nucleus (44,61,62). Such the membrane-bound Nfe2l1 factor can also be activated in the unfolded protein response to the ER stress stimulated by tunicamycin (to inhibit its N-linked glycosylation) (38), in addition to its proteasomal ‘bounce-back’ response (63,64). By contrast, the water-soluble Nfe2l2 is segregated by Keap1 within the cytoplamic compartments, where it is targeted for its ubiquintination-mediated proteasomal degradation (40,53). Upon stimulation of Nfe2l2 by oxidative stress, this CNC-bZIP factor is dissociated from the cytoplamic Keap1, so that it is translocated into the nucleus and transactivates ARE-driven target genes. Collectively, these distinctions of Nfe2l1 from Nfe2l2 demonstrate disparate selection of their tempo-spatial activation or inactivation insomuch as to finely tune distinct subsets of target genes. As such, this also presages synergism and antagonism of between Nfe2l1 and Nfe2l2 in regulating distinct cognate gene expression.

In fact, mouse *Nfe2l1*^*–/–*^ embryonic lethality occurred at middle to late gestation starting at E13.5 (65), whereas the double knockout of *Nfe2l11*^*–/–*^*:Nfe2l21*^*–/–*^ caused an earlier death of the mouse embryos at E10.5 due to extensively increased apoptosis and growth retardation induced by severe endogenous oxidative stress (66). The elevated ROS levels resulted from severely impaired expression of antioxidant defense genes (e.g. *Mt-1, Gclm, Gclc, Ferritin H, Ho-1, Nqo1*) in *Nfe2l11*^*–/–*^*:Nfe2l21*^*–/–*^ MEFs, when compared with the individual knockout of *Nfe2l1*^*–/–*^ or *Nfe2l2*^*–/–*^. Hence, it is inferred that Nrf2 is also allowed for a partial compensation for the loss of Nrf1 function in synergistically regulating critical genes for the intracellular redox homeostatic setting at the robust threshold state during embryogenesis. Such partially overlapping functions of both Nfe2l1 and Nfe2l2 are determined by co-expression patterns of the two CNC-bZIP factors (67-69) and their similarity of conserved sequences (37), which are also, though, driven by their respective *ARE*-containing gene promoters (Table S2).

It is, to our surprise, that the evidence presented herein reveals that knockout of *Nfe2l1* in mice leads to significant decreases of both mRNA and protein levels of Nfe2l2 expressed in *Nfe2l1*^*–/–*^ MEFs. By contrast, almost no changes in mRNA epression levels of *Nfe2l1* in *Nfe2l2*^*–/–*^ MEFs, but the loss of *Nfe2l2* results in obvious decreases in abundances of the full-length Nfe2l1α and its processed proteins (i.e. isoform-B, C & D), as accompanied by an increased Nfe2l1β. Collectively, these results demonstrate that Nfe2l1 acts as a dominant positive regulator to monitor basal constitutive expression of Nfe2l2, at least in MEFs (Figure 8A). In turn, it does not hold true, but Nfe2l2 is required for stabilization of Nfe2l1α and its proteolytic processing to yield a mature CNC-bZIP factor. This notion is further substantiated by the evidence obtained from *Keap1*^*–/–*^ MEFs, showing that obvious accumulation of Nfe2l2 is accompanied by significant increases in abundances of Nfe2l1α and its derivates, but basal mRNA expression of *Nfe2l1* remains to be unaffected.

Recently, emerging evidence showed that the regulatory cross-talks of between Nfe2l2 (but not Nfe2l1) with either α-Pal^NRF1^ or GABPα^NRF2^ are involved in the nuclear control of mitochondrial biogenesis and relevant biological functions (51,52,70). In this study, we unravel that knockout of *Nfe2l1* causes significant decreases in mRNA and/or protein levels of α-Pal^NRF1^ and its target genes *TFAM, Ndufv1, Ndub6, COX5a* and *SOD1* in *Nfe2l1*^*–/–*^ MEFs, whereas knockout of *Nfe2l2*^*–/–*^ leads to marked decreases in protein abundances of α-Pal^NRF1^ and TFAM, but almost unaltered mRNA levels *α-Pal*^*NRF1*^and *TFAM* were observed in *Nfe2l2*^*–/–*^ MEFs. These results indicate that Nfe2l1 acts as a dominant regulator to monitor transcriptional expression of α-Pal^NRF1^, TFAM and other target genes in MEFs, whilst stability of both α-Pal^NRF1^, TFAM proteins may be monitored by Nfe2l1-mediated mechanisms. This is supported by another evidence showing that *Keap1*^*–/–*^ MEFs gave rise to evident increases in basal abundances of Nfe2l2, Nfe2l1α and its processed isoforms, as accompanied by obvious increased proteins of α-Pal^NRF1^ and TFAM. However, only modest increases in their mRNA expression levels of α-Pal^NRF1^ and TFAM were examined in *Keap1*^*–/–*^ MEFs, with an exception of significantly reduced mRNA levels of *Nfe2l2*, but not of *Nfe2l1*. This exception implies that transcriptional expression of Nfe2l2 requires for a putative positive feedback mechanism mediated by Keap1, albeit this Nfe2l2 inhibitor negatively regulates stability of this CNC-bZIP protein. Moreover, our further luciferase assays also revealed that both Nfe2l1 and Nfe2l2 contribute to transcriptional expression of distinct ARE-driven *α-Pal*^*NRF1*^ reporters. Conversely, marked decreases in both mRNA and protein levels of Nfe2l1 and Nfe2l2 resulted from heterogeneous deletion of mouse *α-Pal*^*NRF1+/–*^. Overall, these indicate bidirectional cross-talks of Nfe2l1 and Nfe2l2 with α-Pal^NRF1^ at distinct layers to coordinate the extra-nuclear signalling to the nuclear-to-mitochondrial respiratory and antioxidant transcription networks (as modeled in Figure 8A).

### 3.2 Cross-talks between human Nfe2l1^Nrf1^, Nfe2l2^Nrf2^ and α-Pal^NRF1^ to coordinate the nuclear-to-mitochondrial respiratory and antioxidant gene transcription networks

Intriguingly, we found almost no effects of *hNfe2l1*^*–/–*^ or *hNfe2l2*^*–/–*^ on mRNA and protein levels of human α-Pal^NRF1^ in this experimental setting, albeit both CNC-ZIP genes contain at least one consensus α-Pal binding site within their promoters (Table S3). This is also further confirmed by transcriptomics sequencing of *hNfe2l1*^*–/–*^ *+siNfe2l2*, as compared with those obtained from *hNfe2l1*^*–/–*^ or *hNfe2l2*^*–/–*^ cells. Such results are really contrary to those obtained from mouse *Nfe2l1*^*–/–*^ *and Nfe2l2*^*–/–*^ as described above. Conversely, significantly decreases of human Nfe2l1 and Nfe2l2 protein levels were caused by forced expression of ectopic α-Pal^NRF1^. Similarly, transcriptional activity of *hNfe2l1-luc*, rather than *mNfe2l1-luc*, reporters was also markedly diminished by ectopic α-Pal^NRF1^. By contrast, a recovery of human Nfe2l1 and Nfe2l2 from inhibition of α-Pal^NRF1^ was acquired after this respiratory factor was silenced. Notably, silencing of α-Pal^NRF1^ can only cause a modest decrease in its target TFAM expression, implying that this mitochondrial transcription factor is also monitored by other factors (e.g. GABPα^NRF2^ and/or Pitx2) beyond α-Pal^NRF1^. In this study, we also found that TFAM was down-regulated in *hNfe2l1*^*–/–*^ cells (with accumulation of Nfe2l2), and hence up-regulated in *Nfe2l2*^*–/–*^ cells. Our further evidence reveals that TFAM along with α-Pal^NRF1^ are significantly induced by a redox inducer tBHQ, but such inducible expression levels of both factors are almost completely abolished by *hNfe2l1*^*–/–*^. Collectively, these indicate that Nfe2l1 and Nfe2l2 contribute, respectively, to the putative positive and negative regulation of TFAM, though both CNC-ZIP factors are inhibited by α-Pal^NRF1^ through as-yet-unidentified mechanisms (Figure 8B). Accordingly, the positive regulation of TFAM by Nfe2l1 is further corroborated by its consensus ARE3-driven report assays. However, it should also be noted that the opposing effects of Nfe2l2 on the endogenous TFAM and its consensus *ARE3-luc* reporter could depend on distinct ARE3-adjoining promoter contexts.

Importantly, we have presented the evidence that activity of *hNfe2l1-luc* reporter is transactivated by Pitx2 (albeit it serves as a direct upstream regulator of α-Pal^NRF1^ (39)). Further evidence also reveals that endogenous expression of Nfe2l1 (and Nfe2l2) was significantly up-regulated by over-expression of Pitx2, but rather down-regulated by silencing of this homeobox factor. Such Pitx2-directed alternations of Nfe2l1 and Nfe2l2 are also accompanied by corresponding changes of HO-1, GCLM, as well as α-Pal^NRF1^, TFAM and COX5a. In addition to human Nfe2l1, TFAM is another potential target of Pitx2, because its consensus *PitxRE-luc* reporter was transactivated by this homeobox factor. Altogether with our previously-reported evidence (38,47), these demonstrate multiple regulatory across-talks between Nfe2l1, Nfe2l2 and α-Pal^NRF1^, along with Pitx2 and its target TFAM (Figure 8B), are integrated from the extra-nuclear signalling to the nuclear-to-mitochondrial controls of distinct cellular respiratory and antioxidant gene transcription networks. Yet, they differ between the mouse and human.

## 4. Conclusion

Generally, biological functions and physio(patho)logical responses at different levels are determined by a range of regulatory mechanisms from various signaling pathways to multiple transcription factor-mediated gene networks. Such being the case, these transcription factors and other regulatory molecules could be simplified as key modules which cross-talk among them, in order to form a hierarchical network. In this study, Nfe2l1 and Nfe2l2 are embedded in such a complex molecular interaction network, responsible for antioxidant, detoxification and cytoprotective adaptation to distinct physio-pathological stresses during life process. Of note, Nfe2l1 is endowed with unique dispensable biological functions. This fact confers Nfe2l1 to be distinguished from Nfe2l2 at regulating distinct subsets of ARE-driven cognate genes (e.g., *Aldh1a1* and *GSTa1*). But, the inter-regulatory synergism of Nfe2l1 and Nfe2l2 confers both factors to exert their overlapping roles for co-target genes (e.g, *MT-1, GSTp* and *SOD1*) in MEFs. By contrast, apparent synergistic and antagonistic relationships of between human Nfe2l1 and Nfe2l2 are also demonstrated. Nfe2l1 is a dominant repressor whilst Nfe2l2 is positively regulated by Nfe2l1 in HapG2 cells. Such function of Nfe2l1 is considered as a brake to avoid Nfe2l2 overshoot. For this simplification to be valid, there should always be a correlation between Nfe2l1 and Nfe2l2, so that both CNC-bZIP factors along with cognate target genes are finely tuned in order to meet the changing needs of cells under all conditions. In fact, Nfe2l1 and Nfe2l2 cannot be reduced as a simple module alone. Now, the question becomes what is the minimal modular network to reproduce the observed data and why it is so evolved. At least three of them α-Pal^NRF1^, TFAM and Pitx2 should be included, in addition to Nfe2l1 and Nfe2l2. This defines that the nuclear controls of mitochondrial respiration and biogenesis, as well as protein synthesis and degradation, are physiologically integrated with the extra-nuclear redox signalling to antioxidant cytoprotective responses mediated by Nfe2l1 and/or Nfe2l2. However, in view of mutual inter-regulation of between Nfe2l1 and Nfe2l2, it should be taken severe cautions to interpret the experimental results from loss of *Nfe2l1, Nfe2l2* or both.

## 5. Materials and methods

### 5.1 Chemicals, antibodies and other reagents

All chemicals were of the highest quality commercially available. *tert*-Butylhydroquinone (tBHQ) was from Sangon Biotech (Shanghai, China). Specific antibodies against Nfe2l1 were made from our laboratory (57), antibodies against α-Pal^NRF1^ or V5 ectope were from Abcam and Invitrogen, respectively, whilst β-actin and secondary antibodies were from ZSGB-BIO (Beijing, China). The detailed information about other antibodies, as well as all other key reagents and resources used in this study, was all shown in Table S1.

### 5.2 Expression constructs, Reporter plasmids, and other oligos used for sgRNA or siRNA

Besides four expression constructs for human and mouse Nfe2l1 or Nfe2l2 saved by our group (44,47,62), another four expression constructs for human and mouse Pitx2, and α-pal^NRF1^, were here made by cloning each of those full-length cDNA sequences into the pcDNA3.1 vector. Particularly, the CRISPR/Cas9 plasmid constructs containing guide RNAs specifically targeting mouse α-pal^NRF1^ were created to generate a heterogeneous (*α-pal*^*NRF1+/–*^) knockout cell line from MEFs. Such sgRNAs (listed in Table S1) were designed with highly specific targets for precision genomic positions (as shown in **Figure S1**), before being employed in the sgRNA-directed gene-editing of *α-pal*^*NRF1*^.

Furthermore, several specific *cis*-regulatory luciferase reporter plasmids were prepared from cloning the indicated gene promoter regions. The mouse *Nfe2l1* Promoter region (−2047 to +1210) was amplified by PCR from its genomic loci and then inserted into the pGL3-Basic vector. Another group of the *cis*-regulatory consensus, e.g., ARE and PitxRE (*Pitx2* response elements)-adjoining sequences was cloned from the indicated gene promoter regions, such as *Nfe2l1, TFAM*, and *α-pal*^*NRF1*^ and then inserted into the pGL3-Promoter vector. In addition to those intact reporter genes, such as *PitxRE*-Luc and *ARE*-Luc, relevant point-mutant reporters were engineered here. The fidelity of the above-described constructs was all confirmed to be true by sequencing. All these primers in the above gene manufacture, and other oligos for siRNAs-mediated knockdown of the indicated genes, were listed in Table S1, which all were synthesized by Tsingke (Chengdu, China).

### 5.3 Cell lines, culture and transfection

Wild-type mouse embryonic fibroblasts (MEFs), were given as a gift by Akira Kobayashi. Their relevant knockout MEF lines, e.g., *Nfe2l1*^*–/–*^, *Nfe2l2*^*–/–*^ and *Keap1*^*–/–*^ were also obtained from the groups of Profs. Kobayashi and Hayes. Of note, these cell lines were originally prepared from Prof. Yamamoto‘s laboratory. But, another heterogeneous knockout line of *α-pal*^*NRF1+/–*^ in mice was established by gene-editing in our own laboratory as described below. Besides, the human wild-type (i.e., *hNfe2l1/2*^*+*^*/+*) hepatocellular carcinoma HepG2 cells were originally from the American Type Culture Collection (ATCC, Manassas, VA, USA). The fidelity was conformed to be true by its authentication profiling and STR (short tandem repeat) typing map (by Shanghai Biowing Applied Biotechnology Co., Ltd) (38). On this base, *hNfe2l1α*^*–/–*^ and *hNfe2l2*^*–/–ΔTA*^ were established and characterized in our own laboratory (47). These experimental cell lines were maintained for growth in Dulbecco’s modified Eagle’s medium (DMEM) supplemented with 5 mM glutamine, 10% (v/v) fetal bovine serum (FBS), 100 units/mL penicillin-streptomycin, in the 37 °C incubator with 5% CO_2_. Subsequently, the indicated cells were subjected to transfection for 8 h, which was performed by using Lipofectamine 3000 (Invitrogen, Carlsbad, CA, USA), containing different combinations of indicated plasmids, and then allowed for 24-h recovery from transfection in the fresh medium before relevant experimentation.

### 5.4 Real-time qPCR analysis

Experimental cells were subjected to isolation of total RNAs by using the RNA simple Kit (Tiangen Biotech Co., Beijing, China). Then, 500 ng of total RNAs were added in a reverse-transcriptase reaction to generate the first strand of cDNA (with the Revert Aid First Strand Synthesis Kit from Thermo, Waltham, MA, USA). The synthesized cDNA was served as the template for qPCR, in the GoTaq®qPCR Master Mix (from Promega,), before being deactivated at 95°C for 10 min, and then amplified by 40 reaction cycles of the annealing at 95°C for 15 s and then extending at 60°Cfor 30 s. The final melting curve was validated to examine the amplification quality, whereas the mRNA expression level of β-actin served as an optimal internal standard control. All the primers used for qPCR (Table S1) were synthesized by Tsingke (Chengdu, China).

### 5.5 Western Blotting analysis

Experimental cells were harvested in a lysis buffer (0.5% SDS, 0.04 mol/L DTT, pH 7.5), which was supplemented with the protease inhibitor cOmplete Tablets EASYpack. The lysates were denatured immediately at 100°C for 10 min, sonicated sufficiently, and diluted in 3 × loading buffer (187.5 mmol/L Tris-HCl, pH 6.8, 6% SDS, 30% Glycerol, 150 mmol/L DTT, and 0.3% Bromphenol Blue) at 100°C for 5 min. Then, equal amounts of protein extracts were subjected to separation by SDS-PAGE containing 5–12% polyacrylamide, and subsequent visualization by immunoblotting with distinct primary antibodies as indicated (in Table S1). On some occasions, the blotted membranes were stripped for 30 min and then were re-probed with additional primary antibodies. β-actin served as an internal control to verify equal loading of proteins in each of electrophoretic wells.

### 5.6 Distinct cis-regulatory reporter gene assays

Equal numbers (1.4 × 10^5^) of indicated experimental cells were plated into 12-well plates. When allowed for growth to the density reaching 70–80% cell confluence, they were then transfected with Lipofectamine 3000 reagent (Invitrogen). For distinct lengths of the gene promoter studies, the pGL3-basic vector containing 5-kb (−2248 to +2777), 2.7-kb (−2248 to +483) or 3kb (−259 to +2777) of human *Nfe2l1* promoter, or 3.2kb (−2047 to +1210) of mouse *Nfe2l1* promoter, or an empty pGL3-basic vector as a blank control, was co-transfected for 8 h with an indicated expression construct or an empty pcDNA3.1 as a control, plus another internal control pRL-TK reporter. Subsequently, the cells were allowed for a recovery from transfection in a fresh complete medium for 24 h, before they were lysed in a passive lysis buffer (E1910, Promega, Madison, WI, USA) for the dual-luciferase assay. About 20 μl of the supernatant of cells was assayed for the luciferase activity. Both firefly and renilla luciferase activities in each of the same samples were measured by the Dual-Luciferase reporter assay system (Promega).. For the short consensus sequences of the indicated transcription factor-binding sites, the specific driven luciferase plasmids or the paired point-mutants were co-transfected with the internal control pRL-TK, together with an expression construct for Nrf1, Nrf2, Pitx2 or an empty pcDNA3.1. All the resultant data were normalized and calculated as a fold change (mean ± S.D) relative to the activity of the control group (at a given value of 1.0).

### 5.7 Statistical analysis

The statistical significance of changes was determined using the *Student’s* t-test or *Multiple Analysis of Variations* (MANOVA). All the data presented in this study are shown as a fold change (mean ± S.D), each of which represents at least 3 independent experiments undertaken on separate occasions that were each performed in triplicate.

## Data Availability

All data needed to evaluate the conclusions in the paper are present in this publication and/or the Supplementary Imformation that can be found at xx. Additional data related to this paper may also be requested from the authors.

## Author contributions

S.Z. performed the most experiments with help of Y.X., S.H. and L.Q., except that Y.D. did the remaining experiments and repeated part of the work by S.Z. Both S.Z. and Y.D. collected relative data and also made draft of this manuscript with most figures and supplemental tables. Y.Z. designed and supervised this study, analyzed all the data, helped to prepare all figures with cartoons, wrote and revised the paper.

## Acknowledgments

We are greatly thankful to both Drs. Lu Qiu (at Zhengzhou University, China) and Yonggang Ren (North Sichuan Medical College, Sichuan, China) for having established those relevant cell lines used herein. The study was supported by the National Natural Science Foundation of China (NSFC, with a key program 91429305 and another project 81872336) awarded to Prof. Yiguo Zhang (at Chongqing University, China). This work is also, in part, funded by Sichuan Department of Science and Technology grant (2019YJ0482) to Dr. Yuancai Xiang (at Southwest Medical University, Sichuan, China).

